# Brevican, Neurocan, Tenascin-C and Tenascin-R Act as Important Regulators of the Interplay between Perineuronal Nets, Synaptic Integrity, Inhibitory Interneurons and Otx2

**DOI:** 10.1101/2022.02.24.481837

**Authors:** Cornelius Mueller-Buehl, Jacqueline Reinhard, Lars Roll, Verian Bader, Konstanze F. Winklhofer, Andreas Faissner

## Abstract

Fast-spiking parvalbumin interneurons are critical for the function of mature cortical inhibitory circuits. Most of these neurons are enwrapped by a specialized extracellular matrix structure (ECM) called perineuronal net (PNN), which can regulate their synaptic input. In this study, we investigated the relationship between PNNs, parvalbumin interneurons and synaptic distribution on these cells in the adult primary visual cortex (V1) of quadruple knockout mice deficient for the ECM molecules brevican, neurocan, tenascin-C and tenascin-R. We used super-resolution structured illumination microscopy (SIM) to analyze PNN structure and associated synapses. Additionally, we examined parvalbumin and calretinin interneuron populations. We observed a reduction in the number of PNN-enwrapped cells and a clear disorganization of the PNN structure in the quadruple knockout V1. This was accompanied by an imbalance of inhibitory and excitatory synapses with a reduction of inhibitory and an increase of excitatory synaptic elements along the PNNs. Also, the number of parvalbumin interneurons was reduced in the quadruple knockout, while calretinin interneurons, which do not wear PNNs did not display differences in number. Interestingly, we found the transcription factor Otx2 homeoprotein positive cell population also reduced. Otx2 is crucial for parvalbumin and PNN maturation and a positive feedback loop between these parameters has been described. Collectively, these data indicate an important role of brevican, neurocan, tenascin-C and tenascin-R in regulating the interplay between PNNs, inhibitory interneurons, synaptic distribution as well as Otx2 in the V1.

## 1. Introduction

The extracellular matrix (ECM) is an assembly of extracellular molecules secreted into the cellular microenvironment (Hynes and Naba, 2012). It can provide structural support to tissues through the formation of a meshwork consisting of protein-protein and protein-proteoglycan interactions (Yue, 2014). Furthermore, ECM components can serve as ligands for receptors and transmit signals involved in cellular processes like proliferation, migration, apoptosis, and differentiation (Hynes, 2009, Padhi and Nain, 2020, Wang and Passaniti, 1999). Although the biochemical composition of the ECM is tissue-specific, the main components are collagens, proteoglycans, and glycoproteins (Frantz et al., 2010, Murphy and Rudge, 1985, Maeda, 2015). In the central nervous system (CNS), a specialized ECM called perineuronal net (PNN) can occur around certain neuron populations. Even though the discovery of PNNs by Camillo Golgi took place at the late 19^th^ century (Celio et al., 1998), their important roles in cortical plasticity, regulation of neural activity and in association with neurodegenerative diseases began to emerge only recently (Devienne et al., 2021, Wang and Fawcett, 2012, Lemarchant et al., 2016, Dzyubenko et al., 2016). PNNs enwrap primarily inhibitory gamma-aminobutyric acid (GABA)ergic interneurons by forming a net-like structure around the soma and proximal dendrites (Lensjo et al., 2017). This arrangement of PNN components around the cell is often described as a honeycomb-shaped ECM structure with holes for synaptic contact. The largest population of neurons enwrapped by PNNs in the brain consists of parvalbumin-expressing neurons (Devienne et al., 2021, Hartig et al., 1992). PNNs show considerable molecular heterogeneity in terms of their composition (Miyata et al., 2018), but the main components of a basic general PNN structure consist of a hyaluronan backbone to attach the ECM to the cell surface (Deepa et al., 2006), the chondroitin sulfate proteoglycans (CSPGs) aggrecan, brevican, neurocan, and versican, which belong to the lectican family (Yamaguchi, 2000), and link proteins. Members of the hyaluronan and proteoglycan link protein (HAPLN) family can connect the lecticans to hyaluronan (Bekku et al., 2012). The trimeric glycoprotein tenascin-R (Tnr) can cross-link lectican CSPGs via the C-terminal G3-domain (Morawski et al., 2014, Lundell et al., 2004).

The emergence of PNNs around subsets of neurons is important for synaptic homeostasis. They restrict synaptic plasticity by maintaining synaptic stability, supporting the ability of neurons to adapt their synapses to inhibitory or excitatory signals, and play a key role in synaptogenesis (Bozzelli et al., 2018, Fawcett et al., 2019, Happel, 2016). In the visual cortex, the maturation of parvalbumin-positive interneurons occurs during the onset of the critical period (Lu et al., 2014). A brief well-defined phase, in which sensory input has a major impact on the wiring of local neuronal circuits and synaptic connections (Levelt and Hubener, 2012). The maturation of parvalbumin interneurons is accompanied by the condensation of the ECM as PNNs around these cells (Takesian and Hensch, 2013). It has been proposed that PNNs support the closure of plasticity by synapse stabilization and prevention of synaptic rearrangements and undertake this task even in adulthood (Pizzorusso et al., 2002). Also, plasticity can be reopened even after the critical period by disrupting the PNNs (Pizzorusso et al., 2002). In rodents, reopening plasticity through PNN digestion can cure amblyopia, indicating the importance of PNNs in visual processing (Pizzorusso et al., 2006). This implies the key role of PNNs in the arrangement of inhibitory circuits and their synaptic integrity in the visual cortex as response to visual sensory input. Another important component of parvalbumin interneuron maturation is the orthodenticle homeobox 2 (Otx2) transcription factor. Otx2 is transferred to the visual cortex from extracortical sources including retina and lateral geniculate nucleus in an experience-dependent manner during postnatal development, and also persists into adulthood (Sugiyama et al., 2008, Bernard and Prochiantz, 2016). The internalization of Otx2 by parvalbumin interneurons occurs after binding of the protein to PNNs. When Otx2 binding is disturbed by PNN-degradation or specific inhibitors, the amount of Otx2 in parvalbumin interneurons is reduced (Beurdeley et al., 2012). In addition, a lower internalization of Otx2 by parvalbumin interneurons results in a reduced parvalbumin expression and PNN assembly (Beurdeley et al., 2012). A positive feedback loop between Otx2 and PNNs has recently been reviewed (Bernard and Prochiantz, 2016).

In the present study, we investigated the interplay between PNN components, synaptic stability, inhibitory interneurons, and Otx2 in the primary visual cortex of quadruple knockout mice. The quadruple knockout mouse, which lacks the four crucial ECM molecules brevican, neurocan, Tnc and Tnr, was first described by Uwe Rauch and colleagues (Rauch et al., 2005). This mouse line was generated because previous generated single knockout lines of brevican, neurocan and both tenascins showed no major anatomical deficits (Zhou et al., 2001, Brakebusch et al., 2002, Forsberg et al., 1996, Weber et al., 1999). In contrast, single knockouts of the other lecticans, versican and aggrecan, were embryonically and perinatally lethal (Mjaatvedt et al., 1998, Watanabe et al., 1994). Since the quadruple knockout appeared to be viable and fertile, it turned out to be a suitable model to study the flexibility of the ECM and the effects of the loss of four crucial ECM molecules and their missing interactions simultaneously. An important feature of these four molecules is a strong connection to PNN structures. Brevican, neurocan and Tnr are direct components of PNNs, and Tnc has also been associated with PNNs as a ligand for a variety of CSPGs (Jakovljevic et al., 2021, Galtrey et al., 2008, Morawski et al., 2014, Stamenkovic et al., 2017, Zimmermann and Dours-Zimmermann, 2008). Accordingly, previous studies by our department showed a diminished PNN structure of quadruple knockout hippocampal neurons *in vitro*, accompanied by an impairment of synapse formation and stability. Synaptic activity in these cells was also disturbed, as shown by reduced IPSCs and EPSCs (Geissler et al., 2013). Furthermore, an increase in excitatory synaptic elements and a reduction in inhibitory synaptic elements was observed in hippocampal quadruple knockout cultures, accompanied by an enhancement in neuronal network activity (Gottschling et al., 2019). To gain a deeper insight into the influence of the ECM on synaptic stability, inhibitory interneurons and Otx2, we examined the visual cortex of quadruple knockout mice in this regard. We analyzed the number of PNN-wearing cells and parvalbumin and calretinin interneurons in cortical regions of wildtype control and quadruple knockout mice, highlighting the primary visual cortex (V1). Also, using structured illumination microscopy (SIM), high resolution analyses of the impact of the quadruple knockout on PNN structure were performed. In addition, the distribution of synaptic elements along the PNNs was investigated. Moreover, the occurrence of Otx2 in the V1 and in the retrosplenial cortex (RSC) of wildtype and quadruple knockout mice was examined.

## 2. Material and Methods

### 2.1 Animals

The mice were kept in the animal facility (Faculty of Biology and Biotechnology, Ruhr University Bochum) under a twelve-hour light-dark cycle with free access to chow and water. For the experiments, male and female 129/Sv wildtype (129S2/SvPasCrl; background mouse strain) and quadruple knockout mice (Rauch et al., 2005) were used at 14-16 weeks of age.

### 2.2 Immunohistochemistry

Brains of 14-16 week-old 129/Sv wildtype and quadruple knockout mice were fixed in 4% paraformaldehyde (PFA), cryoprotected and embedded in Tissue-Tek freezing medium (Thermo Fisher Scientific, Cheshire, UK). For cell counting and basic quantification, brain tissue was sectioned coronally (16 μm; (interaural: 1.10 mm, bregma: −2.70 mm) using a cryostat (CM3050 S, Leica). Sections were blocked in blocking solution containing 3% (v/v) normal goat serum (Dianova, Hamburg, Germany), 1% w/v bovine serum albumin (BSA; Sigma-Aldrich) and 0.5% Triton-X-100 (Sigma-Aldrich) in 1x PBS (phosphate-buffered saline) for one hour at room temperature. Primary antibodies (Table 1) were diluted in blocking solution for 24 h. The sections were then washed three times for 10 minutes (min) with 1x PBS. Afterwards, appropriate secondary antibodies were added and incubated for 2 h. Cell nuclei were detected with TO-PRO-3 (1:400; Thermo Fisher Scientific). To analyze PNN structure and to detect synaptic proteins, free-floating sections (40 μm) were used. For the free-floating staining procedure, tissue sections were incubated in 1x PBS for 20 min and then blocked with blocking solution containing 10% (v/v) normal goat serum (Dianova, Hamburg, Germany), 1% w/v BSA and 0.1% (v/v) Triton-X-100 in 1x PBS for one hour at room temperature. Primary antibodies were diluted in blocking solution and incubated at 4°C for three days. Thereafter, tissue sections were washed three times with 1x PBS for 30 min and then incubated with adequate secondary antibodies for two hours.

**Table 1:** Antibodies for Immunohistochemical Stainings.

| Primary antibody/<br>Lectin | Species,<br>clonality/type | Staining<br>procedure | Dilution | Source (stock<br>no.)/RRID | Secondary<br>antibody | Species | Dilution/source |
| --- | --- | --- | --- | --- | --- | --- | --- |
| Aggrecan | Rabbit,<br>polyclonal,<br>IgG | Glass slides | 1:300 | Merck KGaA;<br>AB_90460 | Anti-rabbit<br>Cy5/<br>Anti-rabbit<br>Cy3 | Goat | 1:400<br>Dianova |
| Parvalbu<br>min | Chicken,<br>polyclonal,<br>IgY | Glass slides | 1:100 | Synaptic<br>Systems<br>GmbH;<br>AB_2619887 | Anti-chicken<br>Cy2 | Goat | 1:400<br>Dianova |
| VGLUT1 | Guinea pig,<br>IgG | Free<br>floating | 1:300 | Synaptic<br>Systems<br>GmbH;<br>AB_887878 | Anti-guinea pig<br>Cy5 | Goat | 1:400 |
| VGAT | Guinea pig,<br>polyclonal,<br>IgG | Free<br>floating | 1:300 | Synaptic<br>Systems<br>GmbH;<br>AB_887873 | Anti-guinea pig<br>Cy5 | Goat | 1:400<br>Dianova |
| PSD95 | Mouse,<br>monoclonal,<br>IgG2a | Free<br>floating | 1:300 | Merck KGaA;<br>AB_1121285 | Anti-mouse<br>Cy2 | Goat | 1:400<br>Dianova |
| Gephyrin | Mouse,<br>monoclonal.<br>IgG1 | Free<br>floating |  | Synaptic<br>Systems<br>GmbH;<br>AB_887717 | Anti-mouse<br>Cy2 | Goat | 1:400<br>Dianova |
| Calretinin | Chicken | Glass slides | 1:200 | Synaptic<br>Systems<br>GmbH;<br>AB_2619909 | Anti-Chicken<br>Cy2 | Goat | 1:400<br>Dianova |
| *Wisteria\**<br>*floribunda*<br>*agglutinin* | Lectin | Free<br>floating/Gl<br>ass slides | 1:200 | Vector<br>Laboratories;<br>AB_2336874 | Streptavidin<br>Cy3<br>Cy2 |  | 1:400<br>Dianova |
| Otx2 | Goat,<br>polyclonal,<br>IgG | Free<br>floating | 1:200 | R&D;<br>AB_2157172 | Anti-goat<br>Cy2 | Donkey | 1:400<br>Dianova |

### 2.3 Fluorescence Stereo Microscopy and Confocal Laser-Scanning Microscopy

For counting of aggrecan, calretinin, parvalbumin and WFA (*Wisteria floribunda* agglutinin) positive cells coronal cortex-slices were recorded with a fluorescence stereo microscope (Axio Zoom.V16, Zeiss, Göttingen, Germany). 1.12 mm x 895.11 μm large areas of the V1 were selected in the left and right cortical hemispheres. The number of positive cells was then counted using the ImageJ software (ImageJ 1.51w, National Institutes of Health; Bethesda, MD, USA). The overall aggrecan immunoreactive area [%] was analyzed as previously described (Reinehr et al., 2016, Reinehr et al., 2018). Therefore, images were converted into gray scale and then background subtraction was performed with a rolling radius = 30. Next, lower and upper threshold values were determined for each image. The mean (lower threshold = 5.43 and upper threshold = 70.26) was then used to analyze the percentage of the area fraction coherent. Otx2 staining in the RSC and V1 was examined with a confocal laser-scanning microscope (LSM 510 META; Zeiss, Göttingen, Germany). Two images per animal (630x magnification) were taken. All values were transferred to Statistica software (V13.3; StatSoft (Europe), Hamburg, Germany).

### 2.4 Visualization of PNN Structure

To analyze the PNN structure in wildtype and knockout mice, immunohistochemical stainings of WFA were recorded using fluorescence super-resolution structured illumination microscopy (SIM) on a Zeiss Elyra PS.1 plus LSM880 microscope (Carl Zeiss Microscopy GmbH, Germany). Z-stack imaging with a 63x oil immersion objective (Plan-Apochromat 63x/NA 1.4 OIL DIC, Carl Zeiss Microscopy GmbH, Germany) was used to acquire a stack of 60 slices with an interval of 0.3 μm. During all measurements, laser power and gain were kept constant. For 3D surface rendering and quantitative analysis of PNN parameters, SIM images were imported into IMARIS 9.3.1 (Bitplane AG, Zurich, Switzerland). First, the “*create surface*” tool was utilized to manually draw a surface containing the WFA positive PNN for each optical section. An appropriate threshold was chosen to exclude background signal. A 3D surface containing a single PNN-enwrapped neuron was generated and defined as region of interest (ROI). WFA positive signal outside the ROI was suppressed. The volume of the generated ROI reflects the volume of the PNN. To determine the density of the PNN, a 3D surface of the WFA positive signal inside the ROI was generated and its volume was automatically measured. By dividing the isolated WFA positive volume by the overall PNN volume, the amount of WFA positive signal around the neuron was identified and represented in percentage. Also, the WFA total fluorescence intensity of every PNN was measured via IMARIS and the intensity values were then normalized to the mean intensity value of the wildtype. For each group, 32 PNN-enwrapped cells were analyzed.

### 2.5 Visualization of PNN-associated Synaptic Puncta

Organization of inhibitory and excitatory synapses along PNNs in wildtype and knockout V1 was analyzed. Thus, immunohistochemical stainings either with WFA and antibodies against VGAT and gephyrin or WFA and antibodies against VGLUT1 and PSD95 were performed. Microscopy and generation of 3D surfaces of WFA positive PNNs was accomplished as described in **2.4**. In this ROI, the pre- and postsynaptic puncta distribution at the PNN was analyzed. Therefore, the “spots” analysis tool from the IMARIS software was used. To avoid that background signal was detected as synaptic component, a minimum diameter for synaptic puncta had to be defined. The diameters were determined by measuring the smallest synaptic puncta with the line tool in “slice view”. This resulted in 0.28 μm as diameter for presynaptic markers (VGAT and VGLUT1) and 0.5 μm for postsynaptic markers (gephyrin and PSD95). Next, localization of the spots was identified. “Colocalize spots” was selected and pre- and postsynaptic markers which were within a threshold distance of 1 μm of each other were labeled as colocalized. Spots outside a proximity of 1 μm were labeled as non-colocalized. Again, the synaptic distribution on 32 PNN-enwrapped cells was analyzed for the wildtype and quadruple knockout group.

### 2.6 RNA Purification, cDNA Synthesis and RT-qPCR

Primary visual cortex tissue was dissected and stored at −80°C (n = 7). RNA was isolated using the Gene Elute Mammalian Total RNA Miniprep Kit according to the manufacturer’s protocol (Sigma–Aldrich, St. Louis, MO, USA). Concentration of the isolated RNA was determined photometrically with a BioSpectrometer^®^ (Eppendorf, Hamburg, Germany). cDNA synthesis was performed using a cDNA synthesis kit (ThermoFisher Scientific, Waltham, MA, USA). Therefore, 1 μg RNA was reverse-transcribed with random hexamer primers. For quantitative real-time PCR (RT-qPCR) analyses, the Light Cycler 96^®^ System and SYBR Green I (Roche Applied Science, Mannheim, Germany) was used. Efficiency of the primer pairs (Table 2) was determined via a dilution series of 5, 25 and 125 ng cDNA. For normalization, the housekeeping gene *β-actin* was used.

**Table 2:**
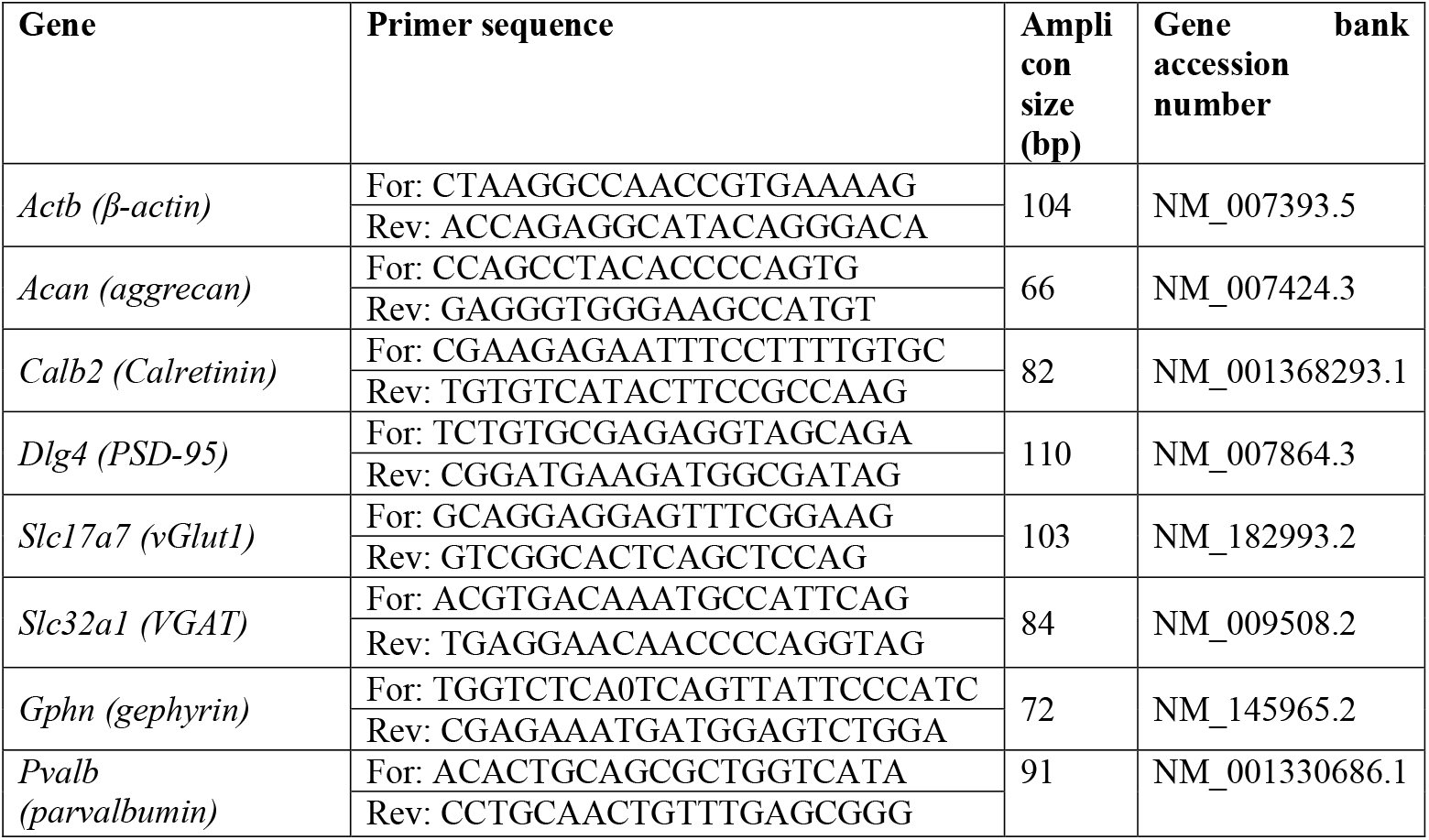
Primer Sequences for RT-qPCR.

### 2.7 Western Blot Analyses

V1 tissue (N = 8/group) was homogenized in 100 μl lysis buffer (60 mM n-octyl-β-D-glucopyranoside, 50 mM sodium acetate, 50 mM tris chloride, pH 8.0 and 2 M urea) supplemented with protease inhibitor cocktail (Sigma-Aldrich) on ice for 1 h. Afterwards, the samples were centrifuged at 14.000 x g at 4°C for 30 min. Then, the protein concentration in the supernatant was determined with a BCA Protein Assay kit (Pierce, Thermo Fisher Scientific, Rockford, IL, USA). To each protein sample (20 μg), 4x SDS was added. Next, samples were denaturized at 94°C for 5 min. Proteins were then separated via SDS-PAGE (10% or 15% gels, respectively, 4-12% polyacrylamide gradient gels). Subsequently, proteins were transferred to a polyvinylidene difluoride (PVDF) membrane (Roth, Karlsruhe, Germany) by Western blotting (1-2 h and 75 mA). Blocking of the membranes was achieved with milk powder (5% w/v milk powder in 1x tris-buffered saline (1x TBS). Additionally, membranes were incubated with primary antibody (Table 3) in blocking solution over night at 4°C. The following day, membranes were washed 3 times for 10 min with 1x TBST (1x TBS with 0.05% Tween^®^20). Incubation with horseradish peroxidase (HRP) coupled secondary antibody (Table 3) in blocking solution for 1h was accomplished at room temperature. Membranes were washed three times with 1x TBST and two times with 1x TBS. For signal detection, ECL substrate solution (Bio-Rad Laboratories GmbH, München, Germany) was applied to the membrane and immunoreactivity was recorded with a MicroChemi Chemiluminescence Reader (Biostep, Burkhardtsdorf, Germany). For the evaluation of the protein signals, band intensity was analyzed using ImageJ software and normalized to a corresponding reference protein (actin). The normalized values of the Western blot results were given in arbitrary units (a.u.).

**Table 3:** List of Primary and Secondary Antibodies for Western Blotting.

| **Primary antibody** | **Species, clonality, type** | **Dilution** | **Source (stock no.)/RRID** | **Secondary antibody, species, type** | **Dilution** | **Source** | **kDa** |
| --- | --- | --- | --- | --- | --- | --- | --- |
| Actin | Mouse, monoclonal, IgG | 1:5000 | BD Bioscience, AB_399901 | Anti-mouse, IgG + IgM HRP | 1:5000 | Dianova | 42 |
| Aggrecan | Rabbit, polyclonal, IgG | 1:10000 | Merck KGaA; AB_90460 | Anti-rabbit, IgG HRP | 1:5000 | Dianova | 150 |
| Parvalbumin | Chicken, polyclonal, IgY | 1:5000 | Proteintech Germany GmbH AB_2880541 | Anti-chicken, IgG HRP | 1:5000 | Dianova | 12 |
| VGLUT1 | Guinea pig, IgG | 1:1000 | Synaptic Systems GmbH, RRID: AB_887878 | Anti-guinea pig, IgG HRP | 1:5000 | Jackson ImmunoResearch Labs | 62 |
| VGAT | Guinea pig, polyclonal, IgG | 1:2000 | Synaptic Systems GmbH; AB_887873 | Anti-guinea pig, IgG HRP | 1:5000 | Jackson ImmunoResearch Labs | 46 |
| PSD95 | Mouse, monoclonal, IgG2a | 1:1000 | Merck KGaA AB_1121285 | Anti-mouse, IgG + IgM HRP | 1:5000 | Dianova | 95 |
| Gephyrin | Mouse, monoclonal. IgG1 | 1:5000 | Synaptic Systems GmbH; AB_887717 | Anti-mouse, IgG + IgM HRP | 1:5000 | Dianova | 93 |
| Calretinin | Chicken | 1:1000 | Synaptic Systems GmbH; AB_2619909 | Anti-chicken, IgG HRP | 1:5000 | Dianova | 31 |
| Vinculin | Mouse, monoclonal, IgG1 | 1:200 | Sigma-Aldrich (V 9131) AB_477629 | Anti-mouse, IgG + IgM HRP | 1:5000 | Dianova | 116 |

### 2.8 Statistical Analyses

Data of immunohistological and Western blot analyses were accomplished via Student’s *t*-test and presented as mean ± standard error mean (SEM) ± standard deviation (SD) using Statistica software (V13.3; StatSoft Europe, Hamburg, Germany). RT-qPCR results were evaluated with the pairwise fixed reallocation and randomization test (REST software) and were presented as median ± quartile ± minimum/maximum. Statistical significance is given by the p value: p < 0.05 = *, p < 0.01 = ** and p < 0.001 = ***. The exact number of experimental repetitions is given in the figure legends.

## 3. Results

### 3.1 Reduced Number of PNNs and Ectopic Shift of Aggrecan in Quadruple Knockout Visual Cortex

Coimmunostaining with WFA and an antibody against aggrecan was performed on murine coronal brain slices of 16 week-old wildtype and quadruple knockout mice. WFA and aggrecan are well described as markers for the visualization of PNNs in the CNS (Hartig et al., 1992, Matthews et al., 2002). An area of interest was chosen (white square) where images were taken for further analyses of the V1 (Fig. 1A). In this area images with a higher magnification were captured and WFA-positive and aggrecan-positive cells were counted (Fig. 1B-E). Significantly reduced numbers of WFA-positive (wildtype: 96.75 ± 8.32, knockout: 64.81 ± 7.42; p < 0.001) and aggrecan-positive (wildtype: 92.21 ± 11.54 vs. knockout: 53.14 ± 7.71; p < 0.001) PNN-enwrapped cells were counted in the quadruple knockout V1 (Fig. 1F). Analyses of other cortical areas near the V1 also showed significant reductions in the number of PNN-enwrapped cells (Supplementary figure 1). Interestingly, despite the reduced number of aggrecan-positive PNNs in the quadruple knockout, no differences in the aggrecan mRNA expression could be detected between the wildtype and quadruple knockout V1 (1.3-fold, p = 0.2, Fig. 1G). Furthermore, aggrecan protein levels, detected as a prominent band at 150 kDa (Fig. 1H) by Western blot analyses, were comparable between both genotypes (wildtype: 0.19 ± 0.03 a.u. vs. knockout: 0.19 ± 0.05 a.u., p = 0.98, Fig. 1I). Considering that the number of aggrecan-positive cells was reduced in the knockout, but aggrecan mRNA expression and protein levels did not differ from the wildtype, laser-scanning microscopy with a higher magnification of immunostained aggrecan-positive PNNs was performed to compare the total aggrecan signal and localization (Fig. 1 J-M). The statistical evaluation of the aggrecan positive area showed no differences between the wildtype and knockout V1 (wildtype: 8.55 ± 9.2% aggrecan-positive area vs. knockout: 7.31 ± 6.14% aggrecan-positive area, p = 0.79). It can be noted that the aggrecan signal in the gray scale image of the wildtype was strictly located in the perisynaptic area of the PNN-positive cells (Fig. 1J, blue arrows). In contrast, the gray scale image of the knockout showed a weaker signal at the perisynaptic area, but also an aggrecan-positive signal in the neuropil (Fig. 1M, yellow arrows). These data showed a reduced number of PNNs with no differences in the aggrecan mRNA and protein levels in the quadruple knockout, but also indicate an ectopic shift of aggrecan from the perisynaptic space to the surrounding neuropil.

**Figure 1:**
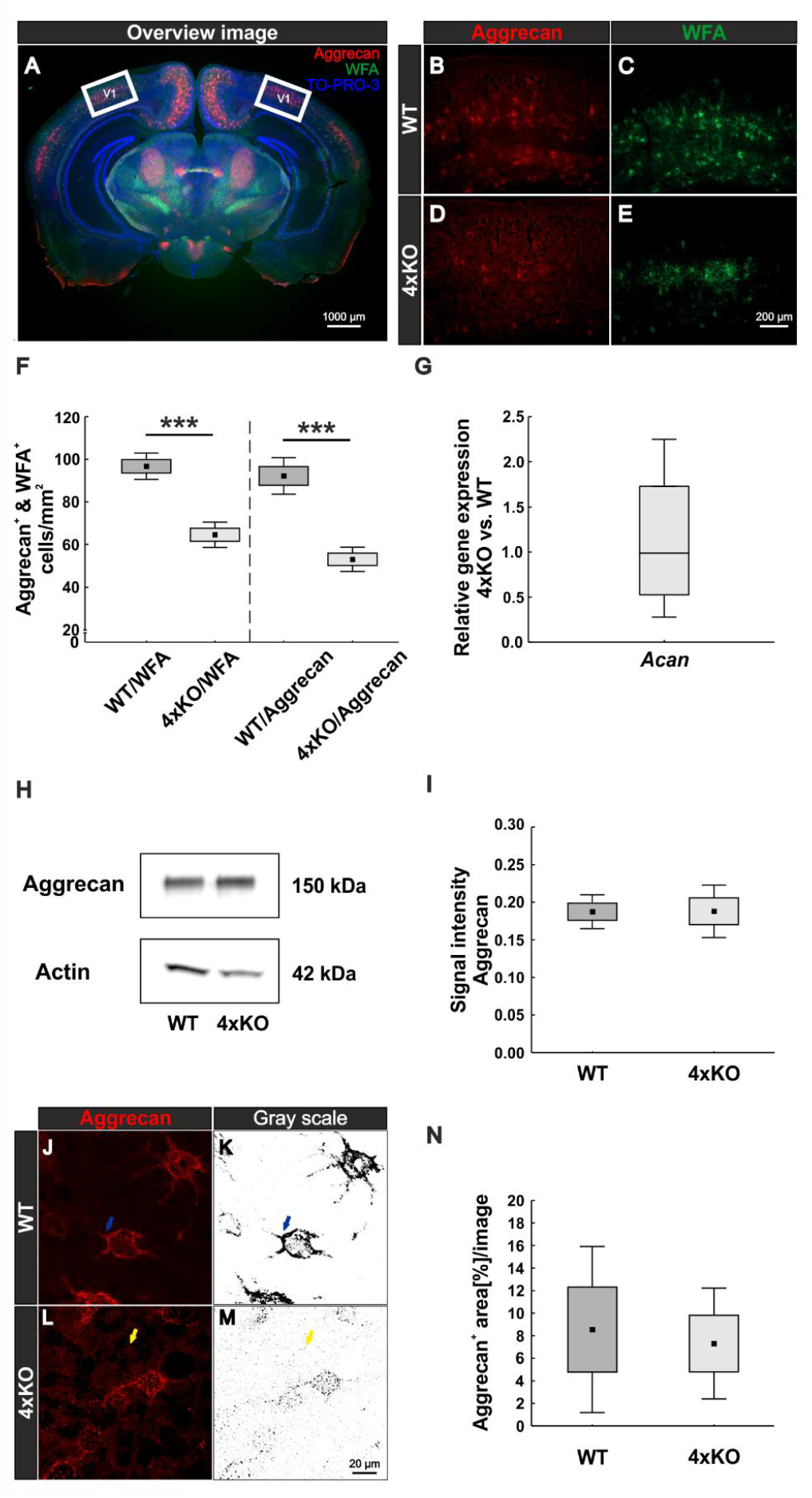
Diminished PNN organization in the visual cortex of quadruple knockout mice. **(A-E)** Immunohistochemical staining of PNNs in murine coronal brain slices with WFA (green) and anti-aggrecan (red). TO-PRO-3 was used as nuclear marker. **(A)** Representative image demonstrating the selected area (white square) for cell counting and further analyses in the visual cortex. **(B-E)** Images of WFA-positive and aggrecan-positive PNN-enwrapped neurons were taken and counted. **(F)** A significantly reduced number of WFA-positive and aggrecan-positive cells (p < 0.001) in the V1 of quadruple knockout mice could be noticed (N = 7). **(G)** RT-qPCR analyses revealed a comparable *Acan* mRNA expression in the visual cortex of wildtype and quadruple knockout mice (p = 0.98, N = 6). **(H)** Western blot analysis of aggrecan protein levels in the V1. **(I)** No differences in the aggrecan protein band intensity were detectable in visual cortex tissue of wildtype and quadruple knockout mice (p = 0.98, N = 8). **(J, L)** For further analyses aggrecan-positive interneurons were documented per laser-scanning microscopy with 630x magnification. **(K, M)** Grey scale images of the analyzed aggrecan-positive signal. Yellow arrows indicate an ectopic shift from the perisynaptic space to the surrounding neuropil in the quadruple knockout mouse. **(N)** Quantification of the stained area showed no differences in the total aggrecan immunoreactivity between wildtype and knockout; 4xKO = quadruple knockout, V1 = primary visual cortex, WFA = *Wisteria floribunda* agglutinin, WT = wildtype, *** = p < 0.001 data are shown as mean ± SEM and SD, scale bar A = 1000 μm, scale bar B-D = 200 μm, scale bar J-M = 20 μm.

### 3.2 Impaired PNN Structure in the V1 of Quadruple Knockout Mice

We showed that the number of PNNs in the quadruple knockout was significantly reduced and aggrecan shifted to the surrounding neuropil. Also, it is well described that brevican, neurocan and Tnr are components of PNNs and Tnc is associated with the PNN structure (Jakovljevic et al., 2021, Galtrey et al., 2008, Morawski et al., 2014, Bekku et al., 2003). Therefore, we investigated the impact of the quadruple knockout on the PNN structure in the V1 with high-resolution SIM. WFA was used as a marker because it showed a more distinct staining pattern than antibodies against aggrecan. PNNs in the wildtype visual cortex appeared with the typical honeycomb structure and accumulation on proximal neurites (Fig. 2A). In contrast, quadruple knockout PNNs exhibited a disrupted structure and less intensely stained proximal neurites (Fig. 2D). Images were then transferred to the IMARIS software for further analyses. A 3D surface of individual PNNs was generated and determined as ROI (Fig. 2. B, E). The WFA-positive signal outside the ROI was suppressed. Next, the volume of the ROI was measured via IMARIS, which reflects the volume of the PNN enwrapping an interneuron. We observed a significant reduction in the volume of the quadruple knockout PNNs (wildtype V1: 8161 ± 1213 μm^3^ vs. knockout V1: 6606 ± 1254 μm^3^, p < 0.05, Fig. 2G). Also, the density of the PNNs was examined. Therefore, the “surface tool” of IMARIS was used again to obtain an accurate 3D structure of WFA-positive signal inside of the ROI (Fig. 2C, F). The sum of all volumes of WFA-positive signals at the 3D structure was then automatically measured describing the volume of all WFA-positive PNN components surrounding the neuron. The volume of the WFA-positive PNN components was then divided by the volume of the overall PNN, enwrapping a neuron. Thus, we could identify the amount of WFA-positive PNN components in the overall volume and therefore the density of the PNNs. The wildtype revealed significantly denser PNNs in comparison to the quadruple knockout PNNs, reflecting a reduced distribution of PNN components surrounding the cell (wildtype V1: 10.77 ± 2.79% PNN density vs. knockout V1: 2.88 ± 1.66%, p < 0.05, Fig. 2H). Also, the quadruple knockout PNNs showed a reduced WFA fluorescence intensity (wildtype: 1.00 ± 0.75% vs. knockout: 0.66 ± 0.35%, p < 0.05, Fig. 2I). In conclusion, the elimination of the four ECM genes not only led to a reduced number of PNNs, but also the structure of the remaining knockout PNNs was severely disrupted.

**Figure 2:**
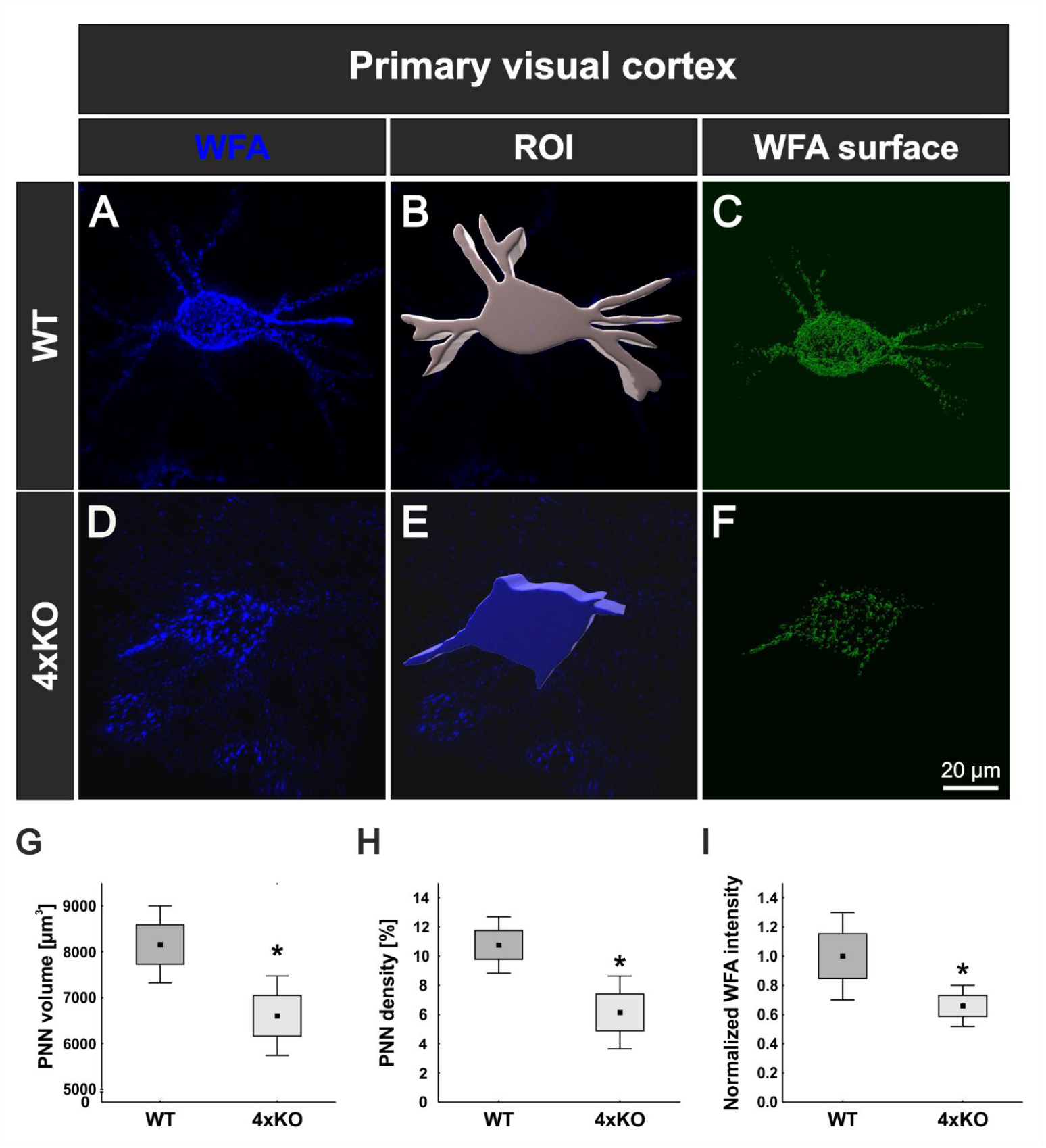
Structural characterization of PNNs in the V1 of wildtype and quadruple knockout mice using super-resolution SIM. **(A, D)** Representative SIM image of WFA-positive PNNs in the wildtype and quadruple knockout V1. As depicted exemplarily, the wildtype PNN showed a typical honeycomb structure with WFA-positive proximal dendrites. In contrast, the quadruple knockout PNN showed a disorganized structure with only one WFA-positive process. **(B, E)** Using IMARIS software, a 3D surface was generated manually containing the WFA-positive signal and the soma enveloped by the PNN, which was determined as ROI. **(C, F)** IMARIS surface technology was then used to create 3D surfaces of the WFA-positive signal inside the ROI best matching PNN anatomy. The WFA signal outside the ROI was suppressed. **(G)** Quantitative analysis revealed a significant reduction in the volume of quadruple knockout PNNs in comparison to wildtype PNNs (p < 0.05). **(H, I)**. Also, PNN density (p < 0.05) and WFA intensity (p < 0.05) were significantly reduced in the quadruple knockout (p < 0.05), indicating fewer WFA-positive PNN components on the cell surface of PNN-enwrapped neurons in the quadruple knockout. 4xKO = quadruple knockout, ROI = region of interest, WFA = *Wisteria floribunda* agglutinin, WT = wildtype, N = 8, * = p < 0.05 data are shown as mean ± standard error mean and standard deviation, scale bar = 20 μm.

### 3.3 Disruption of PNNs Affects Synaptic Integrity

The important role of PNNs in synaptic homeostasis has been well described (Bosiacki et al., 2019, Bukalo et al., 2001, Geissler et al., 2013). PNNs can act as physical barrier to prevent unspecific neuronal connections. Furthermore, PNN components can act as binding partner for inhibitors of synaptic contacts, and they reduce the mobility of ionotropic receptors on the neuronal membrane (Sala et al., 2015, van ‘t Spijker and Kwok, 2017, Saroja et al., 2014). In this context and given that the PNN structure in the quadruple knockout appeared disrupted it was of interest to investigate synapse formation on PNN-enwrapped neurons in wildtype and quadruple knockout mice.

#### 3.3.1 Reduced Number of Structural Inhibitory Synaptic Puncta at Quadruple Knockout PNNs

For the analyses of inhibitory synapses on WFA-positive PNNs, specific antibodies against gephyrin and VGAT were used. Gephyrin and VGAT served as marker for inhibitory postsynaptic and presynaptic elements, respectively, in wildtype and quadruple knockout V1 (Fig. 3A-B’’’). PNNs were singled out as described (Fig. 2) and the synaptic contacts along the PNNs were analyzed (Fig. 3CD’’’). Signals localized outside the ROI were neglected. Synaptic puncta were examined corresponding to their localization. Therefore, spots were generated with the IMARIS software and the tool “Colocalize spots” was used to check the colocalization of pre- and postsynaptic markers (Fig. 3E-F’’). Gephyrin-positive and VGAT-positive puncta which were located within a radius of 1 μm were defined as colocalized. We interpret pre- and postsynaptic puncta in such a spatial proximity as part of a structural synapse (Pyka et al., 2011). Synaptic puncta without a counterpart within a radius of 1 μm were defined as non-colocalized. Most of the gephyrin-positive inhibitory postsynaptic spots seemed to be located at the soma of the cell, whereas inhibitory VGAT-positive presynaptic spots were evenly distributed along soma and proximal dendrites in the wildtype and quadruple knockout (Fig. 3 E, F). Statistical analysis showed that the total number of gephyrin-positive puncta was significantly reduced at the quadruple knockout PNNs in comparison to the wildtype PNNs (wildtype PNNs: 1022. ± 458.39 gephyrin-positive puncta vs. knockout PNNs: 517.32 ± 188.81 gephyrin-positive puncta, p < 0.05, Fig. 3G). In contrast, the total number of VGAT-positive puncta along the PNNs did not differ between the wildtype and quadruple knockout (wildtype PNNs: 3497.91 ± 1204.54 vs knockout PNNs: 2582.86 ± 586.49, p = 0.07, Fig. 3H). However, it is noticeable that the number of VGAT-positive inhibitory presynaptic puncta is clearly larger than the number of gephyrin-positive inhibitory postsynaptic puncta. Colocalized spots were shaded yellow (Fig. 3E’, F’). Here, statistical evaluation showed a significant reduction in the number of colocalized inhibitory gephyrin-positive puncta in the quadruple knockout (wildtype PNNs: 792.54 ± 331.56 gephyrin-positive puncta colocalized vs. knockout PNNs: 340.44 ± 69.12 gephyrin-positive puncta colocalized, p < 0.05, Fig. 3G’). Also, the number of colocalized inhibitory VGAT-positive puncta was significantly reduced at the quadruple knockout PNN in comparison to the wildtype PNN (wildtype PNNs: 930.24 ± 381.09 VGAT-positive puncta colocalized vs. knockout PNNs: 565.9 ± 123.67 VGAT-positive puncta colocalized, p < 0.05, Fig. 3H’). This might be a consequence of the reduced number of gephyrin-positive inhibitory postsynaptic puncta and therefore missing counterparts for the VGAT-positive inhibitory presynaptic puncta at the knockout PNNs. Neither the number of non-colocalized gephyrin-positive puncta (wildtype PNNs: 292.84 ± 170.82 gephyrin-positive puncta non-colocalized vs. knockout PNNs: 176.89 ± 148.00 gephyrin-positive puncta non-colocalized, p = 0.52, Fig. 3G’’), nor the number of non-colocalized VGAT-positive differed between the wildtype and quadruple knockout (wildtype PNNs: 2567.67 ± 977.16 VGAT-positive puncta colocalized vs. knockout PNNs: 2016.67 ± 651.93 VGAT-positive puncta colocalized, p = 0.21, Fig. 3H’). Interestingly, only a small amount of the inhibitory postsynaptic puncta in wildtype and knockout appeared as non-colocalized, whereas a great fraction of the inhibitory presynaptic puncta was non-colocalized. This indicated a more efficient formation of structural postsynapses in comparison to the inhibitory presynapses. Analyses of the protein level and mRNA expression of the inhibitory synaptic marker in V1 tissue showed no differences between wildtype and quadruple knockout (Supplementary figure 2). Therefore, the alteration in the distribution of inhibitory synaptic elements seemed to originate in a disturbed synaptic organization along quadruple PNNs and was not ascribed to an altered number of inhibitory synapses in quadruple knockout tissue. In sum, this result showed a disturbed formation of structural inhibitory synapses at PNNs caused by the combined loss of Tnr, Tnc, brevican and neurocan. Especially the number of inhibitory postsynaptic elements and the formation of structural inhibitory synapses seemed affected.

**Figure 3:**
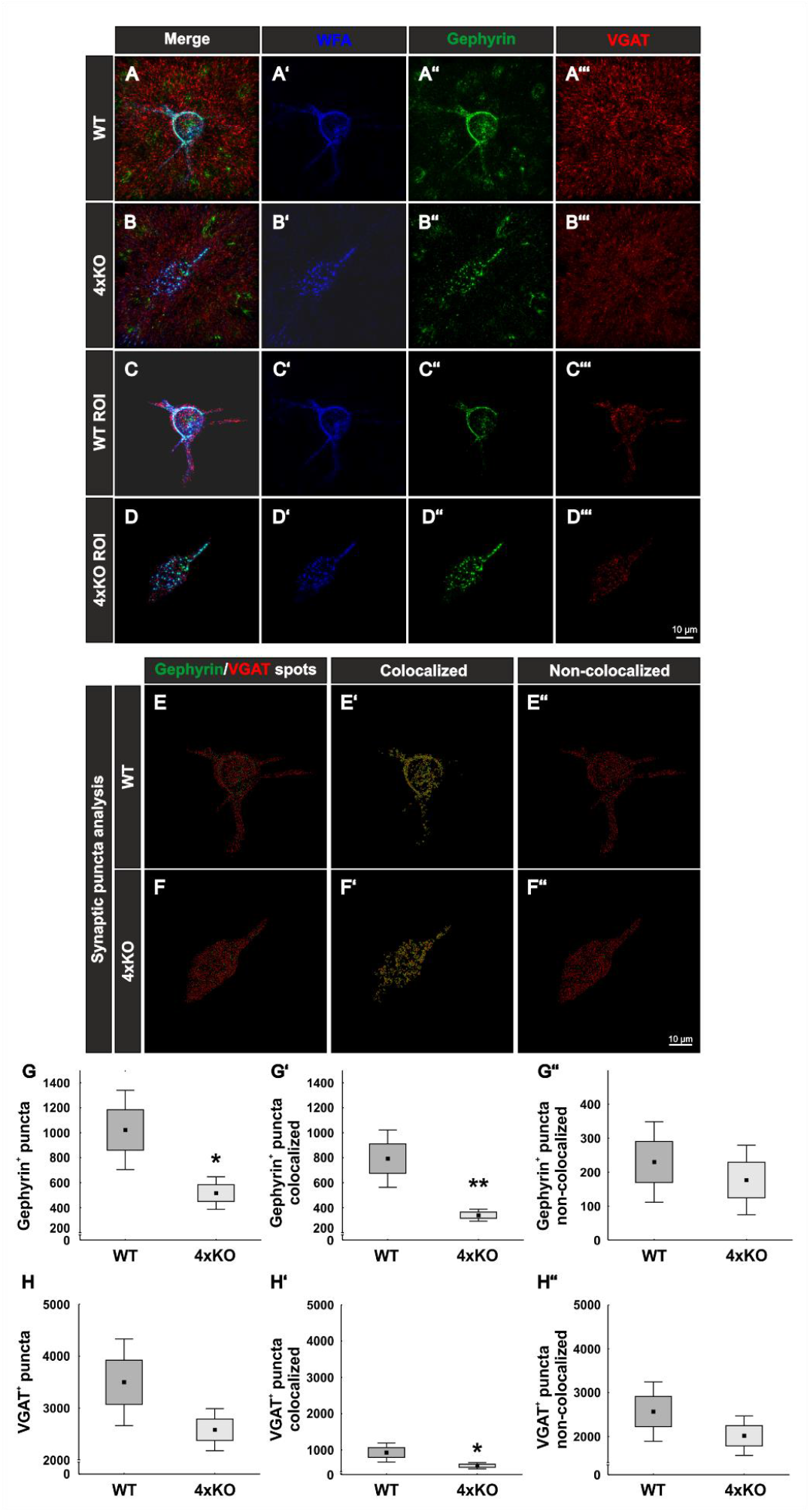
Distribution of inhibitory synaptic elements along PNNs in the V1 of wildtype and quadruple knockout mice. **(A-B’’’)** Immunohistochemical staining of wildtype and quadruple knockout PNNs and inhibitory synapses in the V1. WFA (blue) was used as specific marker for PNNs, antibodies against gephyrin (green) and VGAT (red) as specific markers for inhibitory postsynaptic elements or for inhibitory presynaptic elements, respectively. **(C-D’’’)** A ROI was generated, including solely a single PNN with its perforating inhibitory synapses. **(E-F’’)** Representative image of synaptic puncta represented as spots and checked for their localization via IMARIS. Gephyrin spots are shown in green, VGAT spots in red and colocalized spots are yellow shaded. **(G-G’’)** Statistical evaluation revealed a significant decrease in the total number of gephyrin-positive puncta (p < 0.05) and VGAT and gephyrin double-positive-positive puncta (p < 0.001). No difference in the number of non-colocalized gephyrin-positive puncta (p = 0.52) was detected. **(H-H’’)** Furthermore, the total amount of VGAT-positive puncta was comparable between wildtype and quadruple knockout (p = 0.07), but significantly fewer VGAT-positive and gephyrin-positive colocalized puncta were observed (p < 0.05). The number of non-colocalized VGAT-positive puncta showed no difference between wildtype and quadruple knockout (p = 0.2). 4xKO = quadruple knockout, ROI = region of interest, VGAT = vesicular GABA transporter, WT = wildtype, N = 8; ** = p < 0.01, data are shown as mean ± standard error mean and standard deviation, scale bar = 10 μm.

#### 3.3.2 Increase of Excitatory Synaptic Puncta on PNN-enwrapped Quadruple Knockout Neurons

Next, WFA-positive PNNs in the V1 of wildtype and quadruple knockout mice were examined immunohistochemically for the distribution of excitatory synaptic elements. Therefore, antibodies against PSD95 as a specific marker for excitatory postsynaptic puncta and against VGLUT1 as a specific marker for excitatory presynaptic puncta have been used (Hunt et al., 1996, Naito and Ueda, 1985). To analyze excitatory synaptic elements along wildtype and knockout PNNs, a region of interest containing a single PNN was circumscribed (Fig. 4 A-D’’’). PSD95-positive and VGLUT1-positive signals outside the ROI were suppressed. Next, synaptic spots were generated to analyze the total number of excitatory synapses, but also to evaluate their localization (Fig. 4E-F’’). The distribution of PSD95-positive excitatory postsynaptic puncta appeared unaltered in knockout mice (wildtype PNNs: 371.75 ± 168.29 PSD95-positive synaptic puncta vs. knockout PNNs: 362.05 ± 124.63 PSD95-positive synaptic puncta, p = 0.90, Fig. 4G). The localization of the PSD95-positive synaptic puncta was also similar. No significant differences were found in the number of colocalized puncta (wildtype PNNs: 301.66 ± 148.98 PSD95-positive puncta colocalized vs. knockout PNNs: 262.22 ± 99.11 PSD95-positive puncta colocalized, p = 0.54, Fig. 4G’). Moreover, the number of non-colocalized PSD95-positive puncta was similar between wildtype and knockout PNN-enwrapped neurons (wildtype PNNs: 70.09 ± 27.61 PSD95-positive puncta non-colocalized vs. knockout PNNs: 99.83 ± 39.88 PSD95-positive puncta non-colocalized, p = 0.10, Fig. 4G’’). In contrast, VGLUT1-positive excitatory presynaptic puncta showed an altered distribution along the quadruple knockout PNNs. The number of VGLUT1-positive puncta was significantly increased on PNN-enwrapped neurons of the quadruple knockout compared to the wildtype (wildtype PNNs: 3417.98 ± 784.54 VGLUT1-positive puncta vs. knockout PNNs: 4903.19 ± 913.61 VGLUT1-positive puncta p < 0.01, Fig. 4H). Interestingly, the number of structural VGLUT1-positive synaptic puncta along the PNNs seemed unaltered (wildtype PNNs: 458.06 ± 139.71 colocalized VGLUT1-positive puncta vs. knockout PNNs: 592.44 ± 162.38 colocalized VGLUT1 positive puncta, p = 0.1, Fig. 4H’). Also, a strong increase in the number of non-colocalized VGLUT1-positive puncta could be observed in association with quadruple knockout PNNs compared to wildtype PNNs (wildtype: 2959.89 ± 789.22 VGLUT1-positive puncta non-colocalized vs. knockout PNNs: 4310.75 ± 850.44 VGLUT1-positive puncta non-colocalized, p < 0.01, Fig. 4H’’). Similar to the inhibitory synaptic markers, analyses of the protein and mRNA expression level of excitatory markers in the V1 tissue were comparable between wildtype and quadruple knockout (Supplementary figure 3). Thus, supporting the impression that the altered synaptic distribution along the PNNs was the consequence of an impaired synaptic organization and not the consequence of an altered number of synapses. In conclusion, these observations led to the assumption that the deletion of the four quadruple PNN constituents resulted in an increase of excitatory synaptic elements along the PNN. However, this increase appeared restricted to delocalized excitatory presynapses, whereas excitatory postsynapses and colocalized excitatory presynapses appeared unchanged, indicating a regular distribution of structural excitatory synapses on quadruple knockout PNNs.

**Figure 4:**
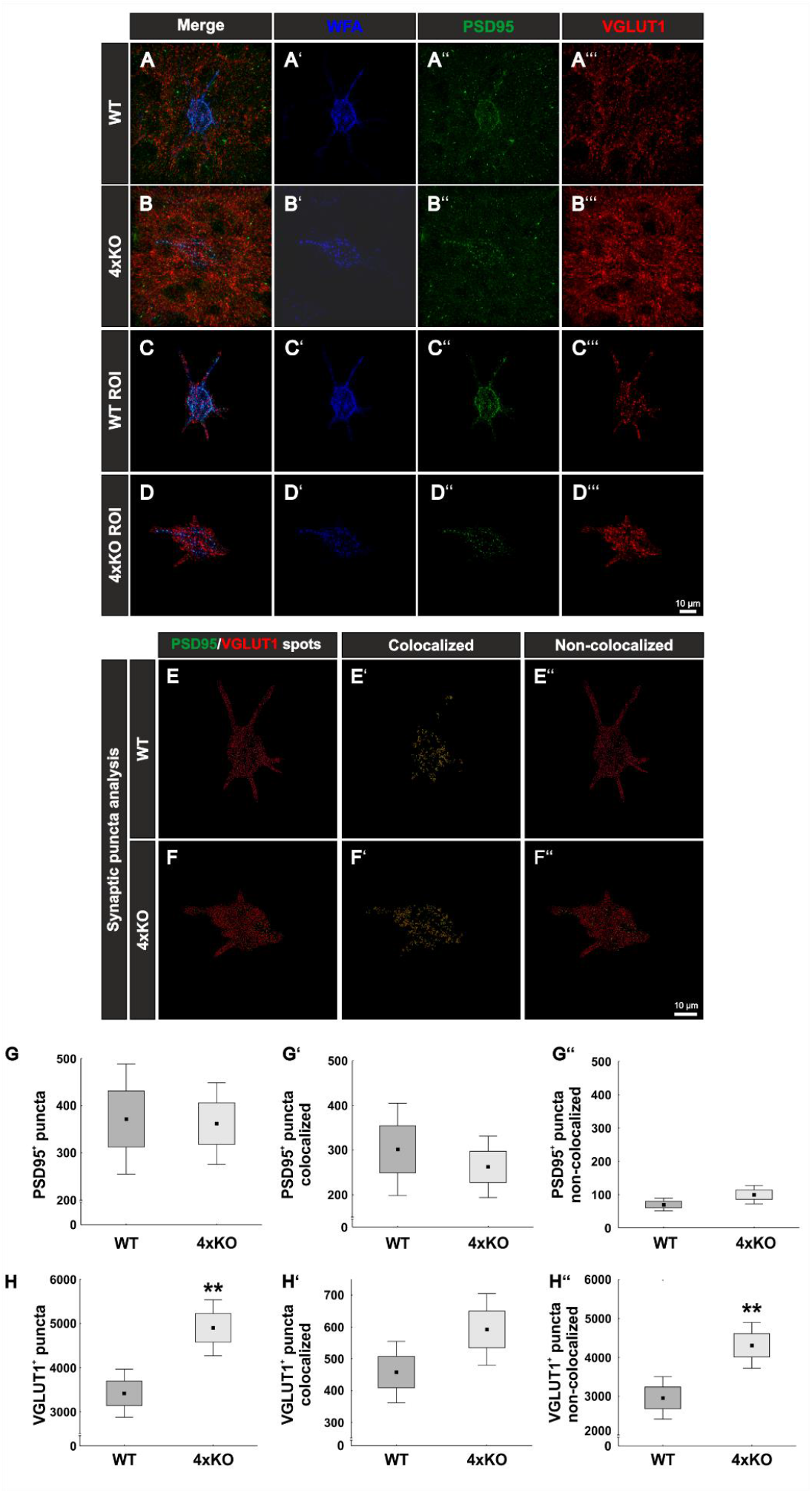
Expression of PSD95-positive and VGLUT1-positive synaptic puncta at wildtype and quadruple knockout PNNs in the V1. **(A-B’’’)** Representative SIM image representing immunohistochemical stainings of WFA-positive PNNs (blue) and excitatory synapses with PSD95 as marker for excitatory postsynapses (green) and VGLUT1 as marker for excitatory presynapses (red). **(C-D’’’)** ROI including single wildtype and quadruple knockout PNNs with its perforating excitatory synapses. **(E-F’’)** For quantitative evaluation, PSD95-positive and VGLUT1-positive synaptic puncta are represented as green and red spots via IMARIS. Colocalized spots are yellow shaded. **(G-G’’)** The total number of PSD95-positive synaptic puncta, the number of PSD95-positive and VGLUT1-positive colocalized synaptic puncta as well as the number of non-colocalized PSD95-positive synaptic puncta along the PNNs was quantified. No difference in the total number of PSD95-positive synaptic puncta (p = 0.90) nor in the number of colocalized (p = 0.54) or non-colocalized PSD95-positive synaptic puncta (p = 0.1) between wildtype and quadruple knockout PNN was detected. **(H-H’’’)** In contrast, the total number of excitatory presynaptic VGLUT1-positive synaptic puncta was significantly increased on the quadruple knockout PNN (p < 0.01). The number of colocalized VGLUT1-positive was similar in the wildtype and quadruple knockout (p = 0.1). However, non-colocalized VGLUT1-positive synapses were significantly reduced on quadruple knockout PNNs (p < 0.01). 4xKO = quadruple knockout, PSD95 = postsynaptic density protein 95, ROI = region of interest, VGLUT1 = vesicular glutamate transporter 1, WT = wildtype, N = 8; ** = p < 0.01, data are shown as mean ± standard error mean and standard deviation, scale bar = 10 μm.

### 3.4 Loss of Parvalbumin-positive Interneurons in the Quadruple knockout Mice

As a next step, it appeared of interest to analyze if the disruption of the PNNs and the synaptic modification along the quadruple knockout PNNs affects the cell population enwrapped in PNNs. In the cortex, PNNs mainly enwrap fast-spiking parvalbumin-positive interneurons (Hartig et al., 1992, Dityatev et al., 2007). To visualize these interneurons, we immunohistochemically double-stained WFA-positive PNNs with antibodies against parvalbumin in wildtype and quadruple knockout V1 (Fig. 5A-F). Most of the parvalbumin-positive interneurons wear PNNs, as indicated by the white arrows. The statistical evaluation showed a significant decrease in the number of parvalbumin-positive cells in the V1 of quadruple knockout mice in comparison to the wildtype (wildtype: 75.94 ± 10.07 parvalbumin-positive cells/mm^2^ vs. knockout: 57.37 ± 18.07, p < 0.05, Fig. 5G). mRNA level of parvalbumin in the V1 of quadruple knockout mice were comparable to the wildtype (0.71-fold, p = 0.09, N = 6, Fig. 5H). Also, parvalbumin protein levels, detected as a prominent band at 12kDa, were comparable in the V1 of wildtype and quadruple knockout mice (p = 0.14, N = 7, Fig. 5I-J). Of particular interest is that this reduction in the number of parvalbumin-positive interneurons was limited to the V1 area. Analyses of other cortex areas adjacent to the V1 showed no difference in the number of parvalbumin-positive cells between wildtype and knockout (Supplementary figure 4). Given such a reduction of parvalbumin-positive interneurons we also examined other interneuron populations in the V1 of wildtype and quadruple knockout. Another marker used for classifying interneurons in the visual cortex is the calcium-binding protein calretinin (Barinka and Druga, 2010). But in contrast to parvalbumin-positive interneurons calretinin-positive interneurons are not enwrapped by PNNs. To analyze the number of calretinin-positive interneurons immunohistochemical stainings with specific antibodies against calretinin were performed (Fig. 5K-P). In contrast to parvalbumin-positive interneurons, calretinin-positive interneurons appeared as not enwrapped by PNNs and as most in close proximity to PNN-enwrapped neurons (yellow arrows). In the case of the calretinin-positive interneuron population, no difference in the number of cells was detected in the V1 of both genotypes (wildtype: 22 ± 2.48 calretinin-positive cells/mm^2^ vs knockout: 22.79 ± 5.71 calretinin-positive cells/mm^2^, p = 0.75, Fig. 5Q). Also, in other cerebral cortex areas, calretinin-positive interneuron populations did not differ between wildtype and quadruple knockout (Supplementary figure 5). *Calretinin* mRNA expression in the V1 of wildtype and quadruple knockout mice was also comparable (1.2-fold, p = 0.38, Fig. 5R). Furthermore, similar calretinin protein levels, detected as a prominent band at 31 kDa (Fig. 5S) by Western blot analyses, were found in wildtype and quadruple knockout V1 tissue (wildtype: 0.976 ± 0.21 a.u. vs. knockout: 0.921 ± 0.16 a.u., p = 0.57, Fig. 5S-T). In summary, these results showed that the parvalbumin-positive interneuron population was significantly reduced in the V1 of quadruple knockout mice, whereas the calretinin-positive interneuron population was comparable to the wildtype population. Whether this apparent loss of parvalbumin-positive interneurons is connected to the disrupted structure of the associated PNNs needs further investigations. It is worth noting that the reduced number of parvalbumin-positive cells was restricted to the V1, whereas other cortical areas seemed unaffected.

**Figure 5:**
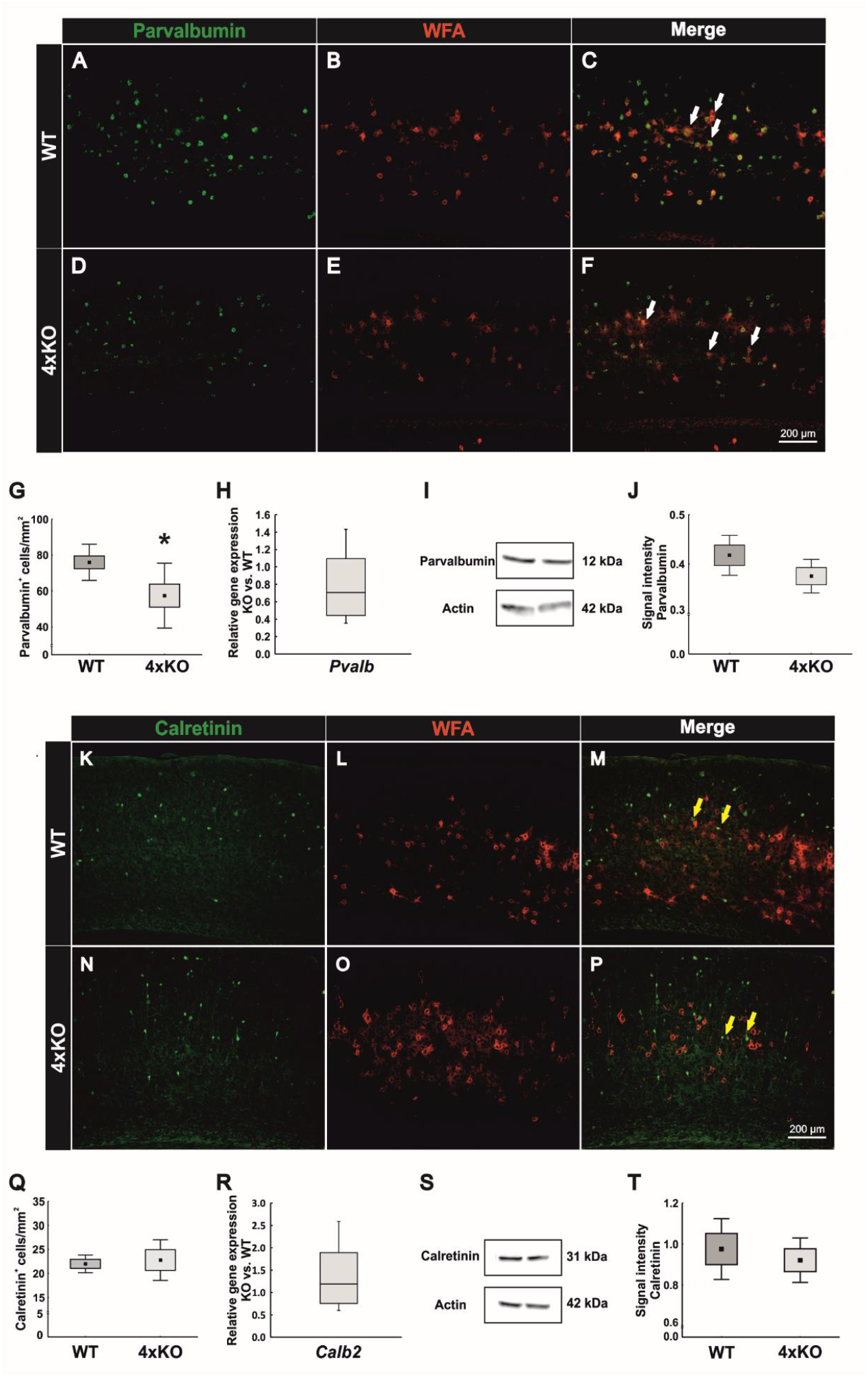
Analyses of parvalbumin-positive and calretinin-positive interneuron populations in the V1 of wildtype and quadruple knockout V1. **(A-F)** Representative coronal V1 brain slices of wildtype and quadruple KO double-labeled using a specific antibody against parvalbumin and WFA in the V1. Parvalbumin-positive fast spiking interneurons were mainly PNN-enveloped (white arrows). **(G)** The number of parvalbumin-positive cells was significantly reduced in the quadruple knockout (p < 0.05, N = 8). **(H)** RT-qPCR analyses revealed a comparable *Pvalb* mRNA expression in the visual cortex of wildtype and quadruple knockout mice (p = 0.09, N = 6). **(I)** Western blot analyses of parvalbumin in visual cortex of wildtype and quadruple knockout. A prominent band was detected at 12 kDa. **(J)** Statistical analyses revealed comparable parvalbumin protein levels between wildtype and quadruple knockout (p = 0.14, N = 7). **(L-P)** Representative coronal V1 brain slices of wildtype and quadruple knockout mice double-labeled with a specific antibody against calretinin and WFA. In contrast to parvalbumin-positive interneurons, calretinin-positive fast spiking interneurons were not enveloped by PNNs (yellow arrows). **(Q)** Statistical analyses showed a comparable number of calretinin-positive interneurons in the wildtype and quadruple knockout V1 (p = 0.75, N = 8). **(R)** Also, *Calb2* mRNA expression levels were comparable between wildtype and quadruple knockout (p = 0.38, N = 6). **(S)** Western blot analyses of calretinin in V1 tissue of wildtype and quadruple knockout. **(T)** No differences in the calretinin protein band intensity were detectable in visual cortex tissue of wildtype and quadruple knockout mice (p = 0.57, N = 8). 4xKO = quadruple knockout, *Calb2* = calbindin 2 (calretinin), *Pvalb* = parvalbumin, WT = wildtype, WFA = *Wisteria floribunda* agglutinin, * = p < 0.05, data are shown as mean ± standard error mean and standard deviation, scale bar = 200 μm.

### 3.5 Lack of Otx2 in the V1 as Possible Trigger for Reduced PV Expression

The remarkable reduction of the parvalbumin-positive interneuron population in the quadruple knockout V1 raised the question about potential causes. In this context it is worth considering the possibility that the selective elimination of parvalbumin-positive interneurons might be causally linked to the structural modification of the associated PNNs in the quadruple knockout mutant. Beyond this option other factors influencing parvalbumin-positive interneuron maturation might be considered. Otx2 is a transcription factor involved in parvalbumin-positive cell maturation. The protein is not synthesized by parvalbumin-positive cells in a cell-autonomous fashion, but rather transferred from other CNS areas, binds with high affinity to PNNs and subsequently accumulates in the interneurons (Beurdeley et al., 2012, Sugiyama et al., 2008). Because of the reported role of PNNs in the trans-neuronal transfer of Otx2, we hypothesized that the structural deficits in the quadruple knockout PNNs should translate in deficits of Otx2 uptake. To test this hypothesis, immunohistochemical double-staining of Otx2 and WFA was performed in coronal brain slices of wildtype and quadruple knockout mice and captured via laser-scanning microscopy (Fig. 6A-M). To determine a possible influence of Otx2 on parvalbumin-positive interneuron populations, two different cortical areas were examined. On the one hand the V1, where a loss of parvalbumin-positive cells in quadruple knockout mice was observed, and on the other hand the RSC where the number of parvalbumin-positive cells was comparable between wildtype and quadruple knockout (Fig. 5 and Supplementary figure 4). The statistical analyses of the Otx2-positive cell number in these areas revealed significant differences. Comparable numbers were seen in the wildtype RSC compared to the quadruple knockout RSC (wildtype RSC: 1305.00 ± 314.37 Otx2-positive cells/mm^2^ vs. knockout RSC: 1250.00 ± 255.33 Otx2-positive cells/mm^2^, p = 0.71, Fig. 6G. In contrast, a significant reduction in the number of Otx2-positive cells was counted in the V1 of quadruple knockout mice compared to the wildtype V1 (wildtype V1: 2140.00 ± 318.89 Otx2-positive cells/mm^2^ vs knockout V1: 1292.50 ± 258.56 Otx2-positive cells/mm^2^, p < 0.001, Fig. 6N). In conclusion, these data showed that the loss of Otx2 was restricted to the V1 in quadruple knockout mice, whereas the RSC seemed unaffected, paralleling the results regarding the parvalbumin-positive cell populations. In the light of these findings, the striking diminution of Otx2 in the V1 of quadruple knockout might represent a plausible cause for the loss of parvalbumin-positive interneurons. It should be noted, that both cortical areas showed a strong disruption of PNNs (Fig. 1 and Supplementary figure 1), which rules out the idea of PNN disruption per se is the origin for parvalbumin-positive and Otx2-positive cell loss in our model. Rather, the reduced capacity of modified PNNs to serve as vehicle for Otx2 transfer presumably compromised the vision-related axonal network where Otx2 plays an important signaling role (Bernard and Prochiantz, 2016).

**Figure 6:**
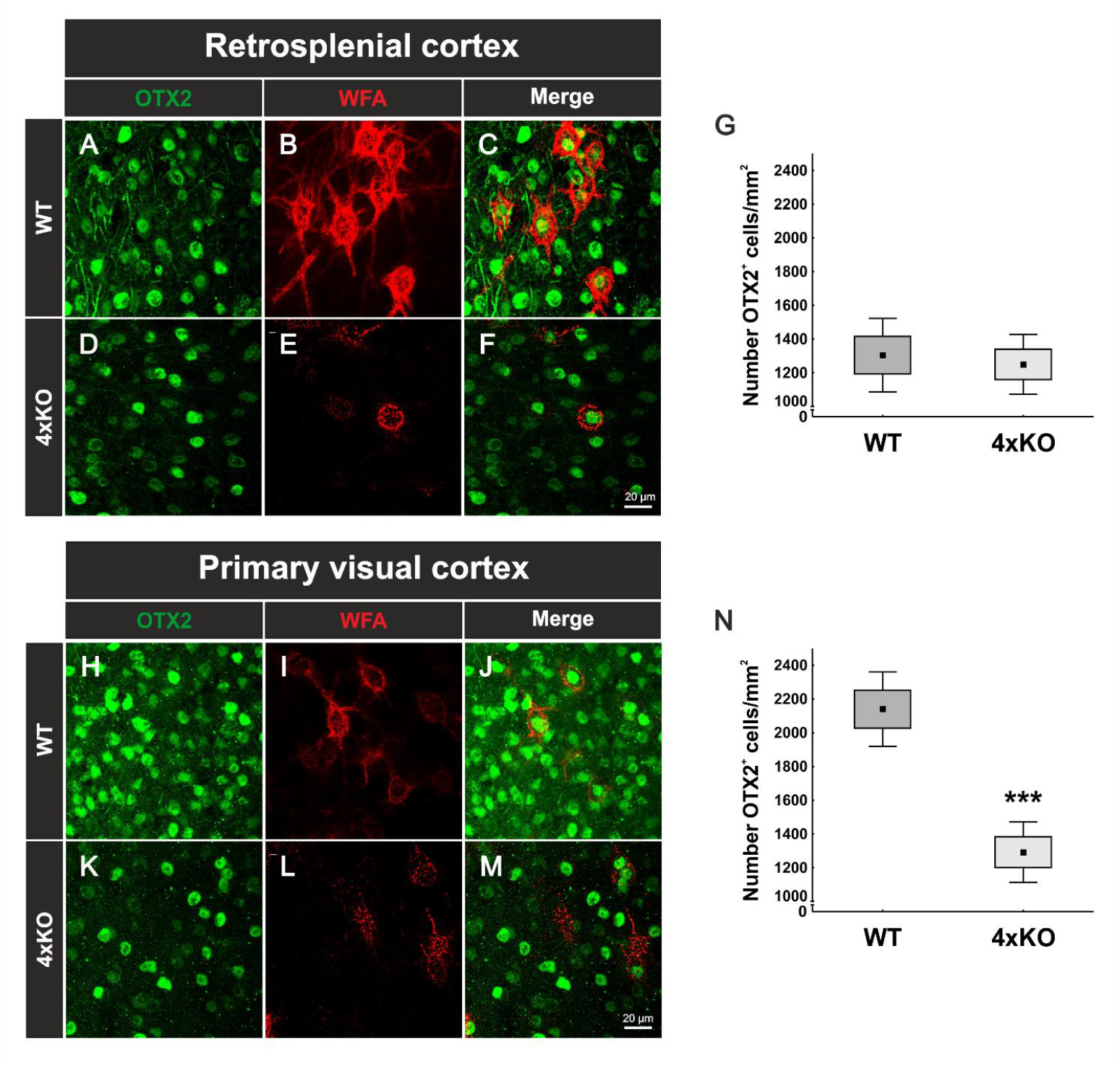
Expression of Otx2 in the cerebral cortex of wildtype and quadruple knockout mice. **(A-M)** Representative images of Otx2-positive cells (green) and WFA-positive PNNs (red) in the RSC and V1 of wildtype and quadruple knockout mice. PNN-enwrapped neurons, but also other cells showed Otx2 accumulation at the soma. **(G)** Counting of Otx2-positive cells in the RSC revealed a comparable number in wildtype and quadruple knockout tissue (p = 0.7, N = 8). **(N)** In contrast, statistical evaluation of Otx2-positive cells in the V1 showed a strong reduction in the number of Otx2-positive cells of quadruple knockout mice in comparison to the wildtype (p < 0.001, N = 8). 4xKO = quadruple knockout, Otx2 = orthodenticle homeobox 2, RSC = retrosplenial cortex, V1 = primary visual cortex, WFA = *Wisteria floribunda* agglutinin, WT = wildtype, *** = p < 0.001 data are shown as mean ± standard error mean and standard deviation, scale bar A = 20 μm.

## 4. Discussion

In this study, we observed a diminished number of PNN-enwrapped cells in the V1 of quadruple knockout mice (Fig. 1F). Also other cerebral cortical areas close to the V1 showed such a reduction in the amount of PNN wearing neurons (Supplementary figure 1). More detailed analyses of the quadruple knockout PNN structure via SIM provided evidence for a strong disruption in the composition of these PNNs (Fig 2 A, D). Single knockout mice lacking either brevican, neurocan, Tnc or Tnr have been described to exhibit altered PNN structures (Faissner et al., 2010). The distribution of carbohydrate epitopes of PNNs on Tnr knockout hippocampal interneurons is abnormal and also a disruption of PNN structure in the cerebral cortex was shown, indicating a disturbance of the molecular scaffolding of ECM components in Tnr knockout PNNs (Weber et al., 1999, Bruckner et al., 2000). Brevican knockout PNNs appeared to be less pronounced and less concentrated near the plasma membrane (Brakebusch et al., 2002). Neurocan knockout mice showed a distinct PNN staining pattern in the olfactory bulb compared to their wildtype littermates and in the cortex an even similar net structure between wildtype and neurocan knockout mice was shown (Hunyadi et al., 2020, Zhou et al., 2001). In fact, Tnc knockout mice have been described as having reduced staining patterns of PNNs compared to wildtype animals, although Tnc is not a direct PNN component and only occurs in close proximity to PNNs (Stamenkovic et al., 2017, Jakovljevic et al., 2021). Compared to the quadruple knockout model, the disturbance of PNN structure in single knockouts appeared to be relatively mild. To our knowledge, no reduction in the number of PNNs is described respectively investigated in the single knockout cortices. This leads to the assumption, that not a single molecule but rather the perturbed interactions between several ECM constituents in the quadruple knockouts cause the intense impairments in structure and number of PNNs. We cannot clarify in this study if the reduced number of PNNs came about by a failure of PNN condensation during development or as consequence a later disorganization of PNNs. An indication for a reduced number of PNNs might be reflected by the observed ectopic shift of aggrecan in the quadruple knockout. Even though the number of aggrecan-positive PNNs was strongly reduced, mRNA expression, protein level and also the immunoreactivity for aggrecan in regard to the overall V1 tissue was similar to WT V1 tissue. This indicates no loss of aggrecan but rather an ectopic shift from the perisynaptic space to the surrounding neuropil of aggrecan in the quadruple knockout V1. It has been shown in mice that the knockout of aggrecan leads to an elimination of PNNs in an *in vitro* cortical culture model as well as in organotypic slice cultures, and, also to an ablated PNN structure *in vivo* (Giamanco et al., 2010, Rowlands et al., 2018). These observations stage aggrecan as the main functional constituent and orchestrator of PNN assembly. Trimeric Tnr can bind three lecticans simultaneously and has been shown to stabilizes PNN formation by clustering aggrecan (Galtrey et al., 2008, Morawski et al., 2014). Furthermore, trimeric Tnr but also hexameric Tnc can cross-link the G3 domains of lecticans and thus tie up the extracellular network (Lundell et al., 2004). Considering the loss of two important scaffolding proteins of the tenascin family, it seems comprehensible that quadruple knockout mice are unable to form a proper PNN structure, which is reflected by the reduced volume, density and WFA intensity of PNNs. It is conceivable that a tampered aggrecan accumulation on PNNs leads to a complete failure of ECM condensation. However, due to the extensive molecular heterogeneity of residual PNNs aggrecan might accumulate trough HAPLN binding and contribute to PNNs with disturbed structure, resulting in a reduced number, but not a total loss of PNNs in the quadruple knockout. These results are consistent with previous *in vitro* studies of hippocampal cultures of quadruple knockout mice, which also revealed a disturbed PNN organization. However, the number of PNN-enwrapped neurons has not been investigated in these cases (Geissler et al., 2013). Another interesting observation in the PNN structure of quadruple knockout mice was the reduced number of PNN-enwrapped neuronal extensions (Supplementary figure 1). Quadruple knockout PNNs tended to surround only one extension, while wildtype PNNs had three to four extensions enwrapped. Since the quadruple knockout had only one PNN-enwrapped extension, it might be the axon, which is the only neuronal extension that has a unique function and appearance compared to the dendrites. In this case, this would indicate a different PNN composition around the axon in comparison to the dendrites, with an important role for the four ECM molecules in the dendritic extracellular environment but not the axonal ECM. However, previous studies have shown a similar composition of the perineuronal micromilieu for diverse cellular domains, contradicting this idea (Bruckner et al., 2006). As previously mentioned, PNNs are an important organizer of synaptic integrity along inhibitory neurons in the cortex (Sigal et al., 2019). *In vivo* studies in the dorsal dentate gyrus of quadruple knockout mice have already shown an altered synaptic function and plasticity (Jansen et al., 2017). Also, hippocampal PNN-enwrapped neurons had a modified number of excitatory and inhibitory synapses, as shown by *in vitro* experiments (Gottschling et al., 2019). In our study, we could show that the number of inhibitory and excitatory synaptic elements as well as their organization is rather different along quadruple knockout PNNs in comparison to wildtype PNNs in the V1. Inhibitory synapses appeared to be more strongly affected than excitatory synapses. In particular, gephyrin-positive inhibitory synaptic puncta were reduced in their number on quadruple knockout PNN-enwrapped neurons. Also, colocalized gephyrin-positive and VGAT-positive puncta were significantly reduced, while non-colocalized inhibitory synapses were unaffected. Apparently, the disruption of the PNNs interferes with the organization of inhibitory synapses. In contrast, PNN disruption in the quadruple knockout V1 did not affect the organization of structural excitatory synapses. The number of structurally colocalized PSD95-positive and VGLUT1-positive synaptic puncta was comparable to wildtype PNNs. However, the number of VGLUT1-positive delocalized synaptic puncta along the quadruple PNN happened to be strongly increased. In conclusion, the synaptic distribution at quadruple knockout PNNs showed an imbalance of excitatory and inhibitory synapses. Similar results were found in *in vitro* studies of hippocampal PNN-enwrapped neurons isolated from quadruple knockout mice. Here, a significantly reduced number of inhibitory and an increased number of excitatory synaptic molecules along the PNNs was described (Gottschling et al., 2019). Also, *in vivo* studies showed that the depletion of PNNs in mature neuronal networks decreases the density of inhibitory synapses (Dzyubenko et al., 2021). mRNA and protein levels of the analyzed synaptic markers in the whole V1 tissue samples, in contrary, showed generally no differences between wildtype and quadruple knockout. This suggests that the altered synaptic organization at the PNNs results from the disruption of its structure and not an effect concerning all the cells in the V1. Organization of synaptic connections by PNNs is well described (Fawcett et al., 2019). They can act as physical barrier to prevent axons of other neurons to connect with the body of its enwrapped neuron (Corvetti and Rossi, 2005, Pyka et al., 2011). An impaired balance between excitation and inhibition contributes to the pathophysiology of autism and related neuropsychiatric disorders (Sohal and Rubenstein, 2019, Nelson and Valakh, 2015, Lee et al., 2017a). Furthermore, interactions between PNNs and parvalbumin interneurons is altered in mouse models of autism (Xia et al., 2021). In this context the quadruple knockout mouse model could be interesting for studies of neurological disorders in the future. The PNN disruption and synaptic alterations in quadruple knockout V1 were accompanied by a reduction in the parvalbumin interneurons. In the visual cortex, PNNs enwrap mainly parvalbumin-positive interneurons (Hartig et al., 1992, Aronitz et al., 2021). In contrast, calretinin interneurons which appeared not to be ensheathed by PNN displayed no alteration in their number of cells in comparison to the wildtype V1. A protective role of PNNs for their parvalbumin neurons is described. In this regard, PNNs can act as a protective shield for parvalbumin interneurons against oxidative stress and potential neurochemical stimuli (Cabungcal et al., 2013, Morawski et al., 2014, Suttkus et al., 2012, Reichelt et al., 2019). Because of their fast-spiking properties parvalbumin interneurons have a high metabolic demand, which renders them sensitive to oxidative stress. Other interneurons such as calretinin-positive interneurons are not particularly sensitive to oxidative stress (Cabungcal et al., 2013). A missing protection by PNNs for parvalbumin interneurons leading to their loss in the quadruple knockout V1 seems plausible, considering the disturbed structure and reduced number of PNNs. And although the occurrence of oxidative stress in this animal model is not documented, glutamate is considered responsible for most of oxidative stress induction in the mammalian brain (Herrera et al., 2001). This scenario is consistent with the increased number of VGLUT1-positive synaptic puncta at quadruple knockout PNNs. In contradiction with this interpretation other cortical areas, where the number of PNNs was reduced as well did not display a reduction in the number of parvalbumin interneurons. Another important factor for parvalbumin maturation is Otx2 (Lee et al., 2017b, Apulei et al., 2019). Otx2 is persistently internalized by parvalbumin-positive interneurons through PNN binding. Also, the disruption of Otx2 localization to parvalbumin interneurons reduces parvalbumin and PNN expression (Beurdeley et al., 2012). Interestingly, similar to parvalbumin interneurons, the number of Otx2-positive cells was reduced in the V1, but not in the RSC of quadruple knockout mice. It appears likely that the reduced number of Otx2-positive cells cells is causally linked to the missing parvalbumin interneurons in the quadruple knockout V1. However, the reduction of Otx2-positive cells appeared considerably more extensive than the reduction in parvalbumin interneurons. It is postulated that transfer of Otx2 from visual pathways to the V1 depends on visual experience. Enucleation of the eyes or dark-rearing from birth reduces Otx2 protein levels, which leads to a loss of Otx2-positive cells, weakens parvalbumin expression, and results in a reduced number of WFA-labeled cells compared to intact animals. Also, calretinin-positive cells were not affected in their number (Sugiyama et al., 2008). Thus, the interplay between PNNs, interneurons and Otx2 in the V1 of quadruple knockout mice seems reminiscent of mouse models with a deprivation in visual experience and input. PNN degradation by digesting CSPGs coupled to visual sensory input altered synapse selectively onto parvalbumin interneurons and showed PNN control of visual processing (Faini et al., 2018). Therefore, brevican, neurocan, Tnc and Tnr seem to be important regulators of visual system integrity, especially by virtue of their role as PNN components. Regarding this data, future studies on the impact of the knockout of the four ECM molecules on visual processing might be interesting for a better understanding of ECM involvement in the organization of visual pathways.

## 5 ETHICS STATEMENT

Animal care and experimental procedures were performed in accordance with the Society for Neuroscience and the EU animal welfare protection laws. The study was supervised by the animal welfare commissioner of the Ruhr University Bochum.

## 6 AUTHORS CONTRIBUTION

CM wrote the manuscript. CM, JR and AF designed the study. CM, JR, LR and VB performed the experiments and analyzed data. JR, LR, VB, KFW and AF revised the manuscript. All authors read and approved the final manuscript.

## 9 ACKNOWLEDGEMENTS

We thank Anja Coenen, Sabine Kindermann, and Franziska Mennes for technical assistance. We thank Dr. D. Wegrzyn for helpful discussions.

## 10 ABBREVIATIONS

BSA: bovine serum albumin
CNS: central nervous system
CSPG: chondroitin sulfate proteoglycan
ECM: extracellular matrix
GABA: gamma-aminobutyric acid
HAPLN: hyaluronan and proteoglycan link protein
KO: knockout
min: minutes
Otx2: orthodenticle homeobox 2
PBS: phosphate-buffered saline
PFA: paraformaldehyde
PNN: perineuronal net
RT-qPCR: quantitative real time PCR
PSD95: postsynaptic density protein 95
ROI: region of interest
RSC: retrosplenial cortex
SD: standard deviation
SEM: standard error mean
SIM: structured illumination microscopy
TBS: tris-buffered saline
Tnc: tenascin-C
Tnr: tenascin-R
V1: primary visual cortex
VGAT: vesicular GABA transporter
VGLUT1: vesicular glutamate transporter 1
WFA: *Wisteria floribunda* agglutinin
WT: wildtype

## 11 CONTRIBUTIONS TO THE FIELD STATEMENT

Perineuronal nets (PNNs) are a specialized form of extracellular matrix surrounding mainly interneuron subpopulations in the central nervous system. Degradation of PNNs affects synaptic integrity and neuronal activity, which indicates the importance of this structure for proper neuronal function. Furthermore, neurological disorders like schizophrenia and Alzheimer’s disease, but also autism spectrum disorder are accompanied by alterations of PNN-enwrapped parvalbumin-expressing interneurons. In our manuscript we analyzed quadruple knockout mice, lacking the four ECM molecules brevican, neurocan, tenascin-C and tenascin-R, as suitable model for investigations on PNN composition and PNN-enwrapped neurons. Referring to super-resolution microscopy analyses on PNN-enwrapped neurons and the distribution of their synaptic elements in this transgenic mouse model, we suggest an important role of brevican, neurocan, tenascin-C and tenascin-R in the regulation of the interplay between PNNs, synaptic integrity, inhibitory interneurons and the transcription factor Otx2. In this regard, the quadruple knockout mouse is presented as a tool for future studies on the influence of the ECM on visual processing. Furthermore, our study offers a novel approach for the investigation of PNN structure in the context of synaptic distribution via super-resolution microscopy and image analyses based on the IMARIS software.

