## Supplementary Material for "Brevican, Neurocan, Tenascin-C and Tenascin-R Act as Important Regulators of the Interplay between Perineuronal Nets, Synaptic Integrity, Inhibitory Interneurons and Otx2"

#### 16 Supplementary FIGURE LEGENDS

**Supplementary Figure 1:** Diminished PNN organization in the retrosplenial-, secondary visual- and auditory cortex of quadruple knockout mice. **(A-F)** Immunohistochemical staining of PNNs in murine coronal brain slices with WFA (green) and anti-aggrecan (red). Images of WFA-positive and aggrecan-positive PNN-enwrapped neurons were taken and counted in the retrosplenial-, secondary visual- and auditory cortex. **(G)** A significantly reduced number of WFA-positive cells in the retrosplenial-, secondary visual- and auditory cortex of quadruple knockout mice could be noticed ($p < 0.001$, N = 7). **(H)** Also, the number of aggrecan-positive cells was significantly reduced in retrosplenial-, secondary visual- and auditory cortex of quadruple Knockout mice ($p < 0.001$, N = 7). **(I, J)** The number of WFA-positive and aggrecan-positive processes per PNN were counted in retrosplenial-, secondary visual-, primary visual- and auditory cortex. Both, WFA-positive and aggrecan-positive processes were significantly reduced in the examined cortical areas ($p < 0.001$, N = 7); 4xKO = quadruple knockout, Aud = auditory cortex, V1 = primary visual cortex, V2 = secondary visual cortex, RSC = retrosplenial cortex, WFA = Wisteria floribunda agglutinin, WT = wildtype, *** = $p < 0.001$ data are shown as mean ± SEM and SD, scale bar = 20 µm.

**Supplementary Figure 2:** Inhibitory synaptic elements in the V1 of quadruple knockout mice. **(A)** Western blot analysis of gephyrin protein levels in the V1. **(B)** No significant differences in the gephyrin protein band intensity were detectable in visual cortex tissue of wildtype and quadruple knockout mice (p = 0.16, N = 8). **(C)** RT-qPCR analyses revealed a comparable *Gephn* mRNA expression in the visual cortex of wildtype and quadruple knockout mice (p = 0.07, N = 6). **(D)** Western blot analysis of gephyrin protein levels in the V1. **(E)** Comparable VGAT protein band intensity in visual cortex tissue of wildtype and quadruple knockout (p = 0.14, N = 8). **(F)** RT-qPCR analyses revealed a significant lower *Slc32a1(VGAT)* mRNA expression in the visual cortex of quadruple knockout mice ($p < 0.001$, N = 6); 4xKO = quadruple knockout, *Gephn* = *Gephyrin*, V1 = primary visual cortex, VGAT = vesicular GABA transporter, WT = wildtype, * = $p < 0.05$ data are shown as mean ± SEM and SD.

**Supplementary Figure 3:** Excitatory synaptic elements in the V1 of quadruple knockout mice. **(A)** Western blot analysis of PSD95 protein levels in the V1. **(B)** No significant differences in the PSD95 protein band intensity were detectable in visual cortex tissue of wildtype and quadruple knockout mice (p = 0.85, N = 8). **(C)** RT-qPCR analyses revealed a comparable *Dlg4* mRNA expression in the visual cortex of wildtype and quadruple knockout mice (p = 0.10, N = 6). **(D)** Western blot analysis of VGLUT1 protein levels in the V1. **(E)** Comparable VGLUT1 protein band intensity in visual cortex tissue of wildtype and quadruple knockout (p = 0.46, N = 8). **(F)** RT-qPCR analyses revealed comparable *Slc17a7 (VGLUT1)* mRNA expression in the visual cortex of wildtype and quadruple knockout mice (p = 0.40, N = 6); 4xKO = quadruple knockout, *Dlg4 = postsynaptic density protein 95*, PSD95 = postsynaptic density protein 95, Slc17a7 = vesicular glutamate transporter 1, V1 = primary visual cortex, VGLUT1 = vesicular glutamate transporter 1, WT = wildtype, * = $p < 0.05$ data are shown as mean ± SEM and SD.

**Supplementary Figure 4:** Analyses of parvalbumin-positive interneuron populations in the retrosplenial-, secondary visual- and auditory cortex of wildtype and quadruple knockout mice. **(A-F)** Representative coronal cortical brain slices of wildtype and quadruple KO double-labeled using a specific antibody against parvalbumin and WFA. **(G)** The number of parvalbumin-positive cells was comparable in the retrosplenial cortex in wildtype and quadruple knockout mice (p = 0.67, N = 8). Furthermore, the number of parvalbumin-positive cells was comparable in the secondary visual cortex (p = 0.11, N = 8) and the auditory cortex of wildtype and quadruple knockout mice (p = 0.61, N = 8).

4xKO = quadruple knockout, *Pvalb* = parvalbumin, WT = wildtype, WFA = *Wisteria floribunda* agglutinin, * = $p < 0.05$, data are shown as mean ± SEM and SD, scale bar = 200 µm.

**Supplementary Figure 5:** Analyses of parvalbumin and calretinin-positive interneuron populations in the retrosplenial-, secondary visual- and auditory cortex of wildtype and quadruple knockout mice. **(A-F)** Representative coronal cortical brain slices of wildtype and quadruple KO double-labeled using a specific antibody against calretinin and WFA retrosplenial-, secondary visual- and auditory cortex of wildtype and quadruple knockout mice. **(G)** The number of calretinin-positive cells was comparable in the retrosplenial cortex between wildtype and quadruple knockout mice ($p = 0.63$, $N = 8$). Furthermore, the number of calretinin-positive cells was comparable in the secondary visual cortex ($p = 0.53$, $N = 8$) and the auditory cortex of wildtype and quadruple knockout mice ($p = 0.19$, $N = 8$). 4xKO = quadruple knockout, *Pvalb* = parvalbumin, WT = wildtype, WFA = *Wisteria floribunda* agglutinin, * = $p < 0.05$, data are shown as mean ± SEM and SD, scale bar = 200 µm.

### 17 SUPPLEMENTARY FIGURES

**Supplementary Figure 1:**

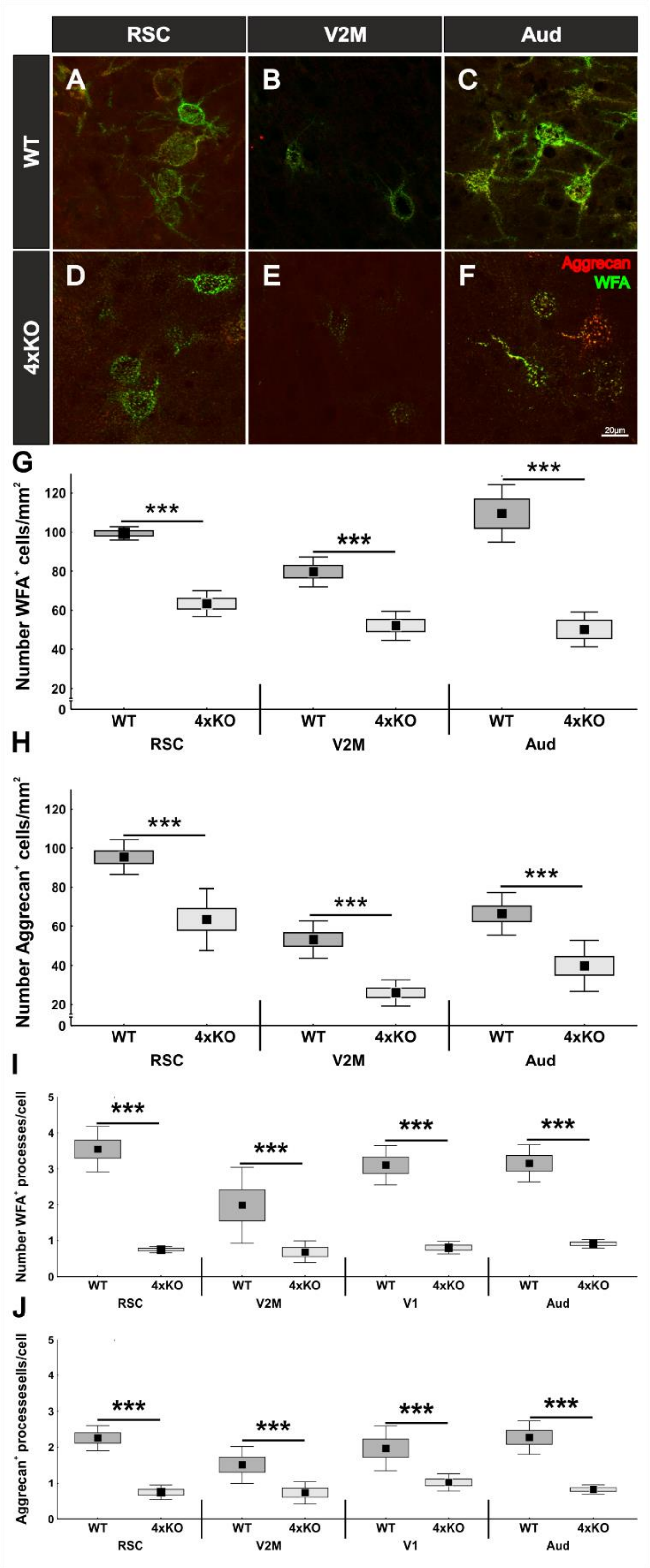

**Supplementary Figure 2:**

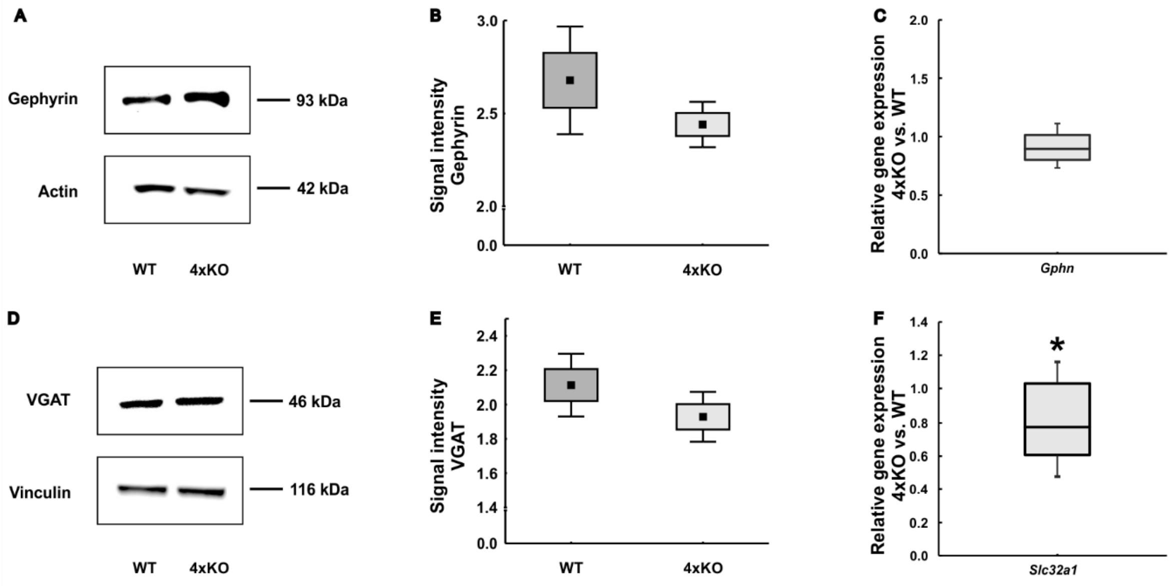

**Supplementary Figure 3:**

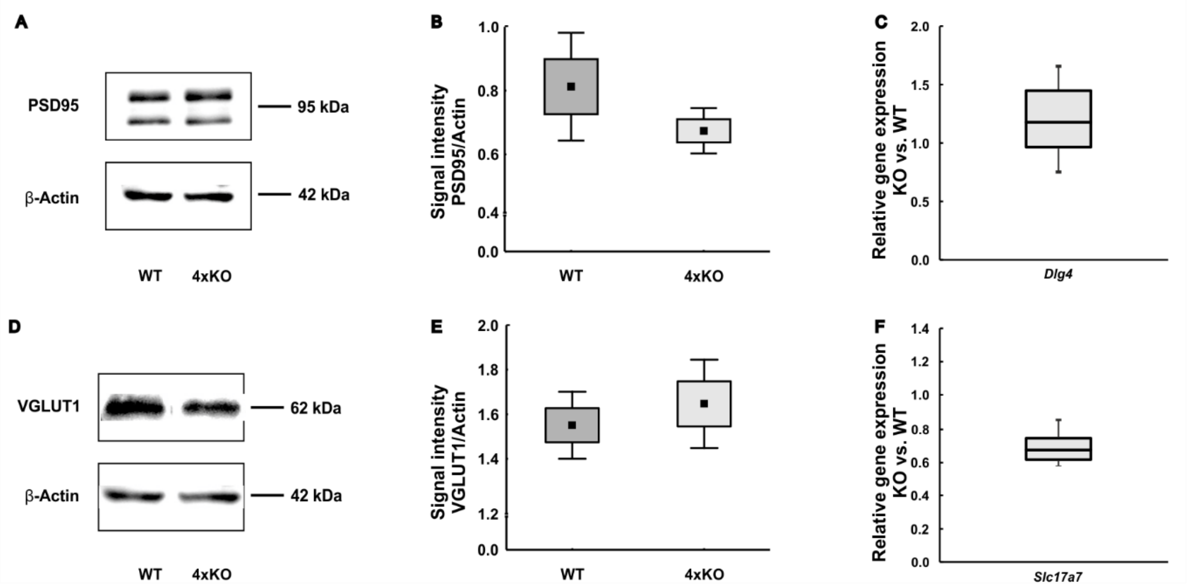

**Supplementary Figure 4:**

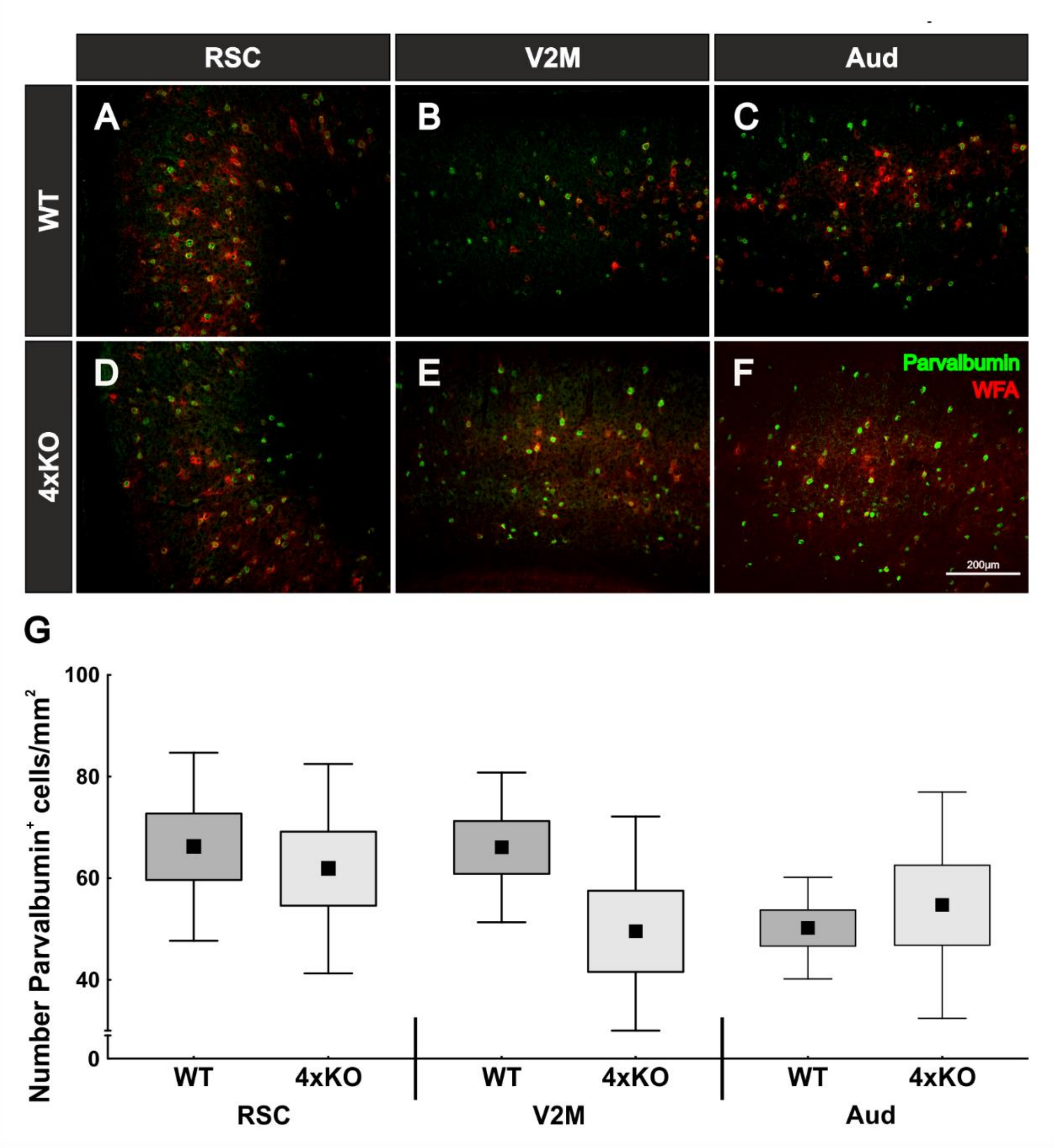

**Supplementary Figure 5:**

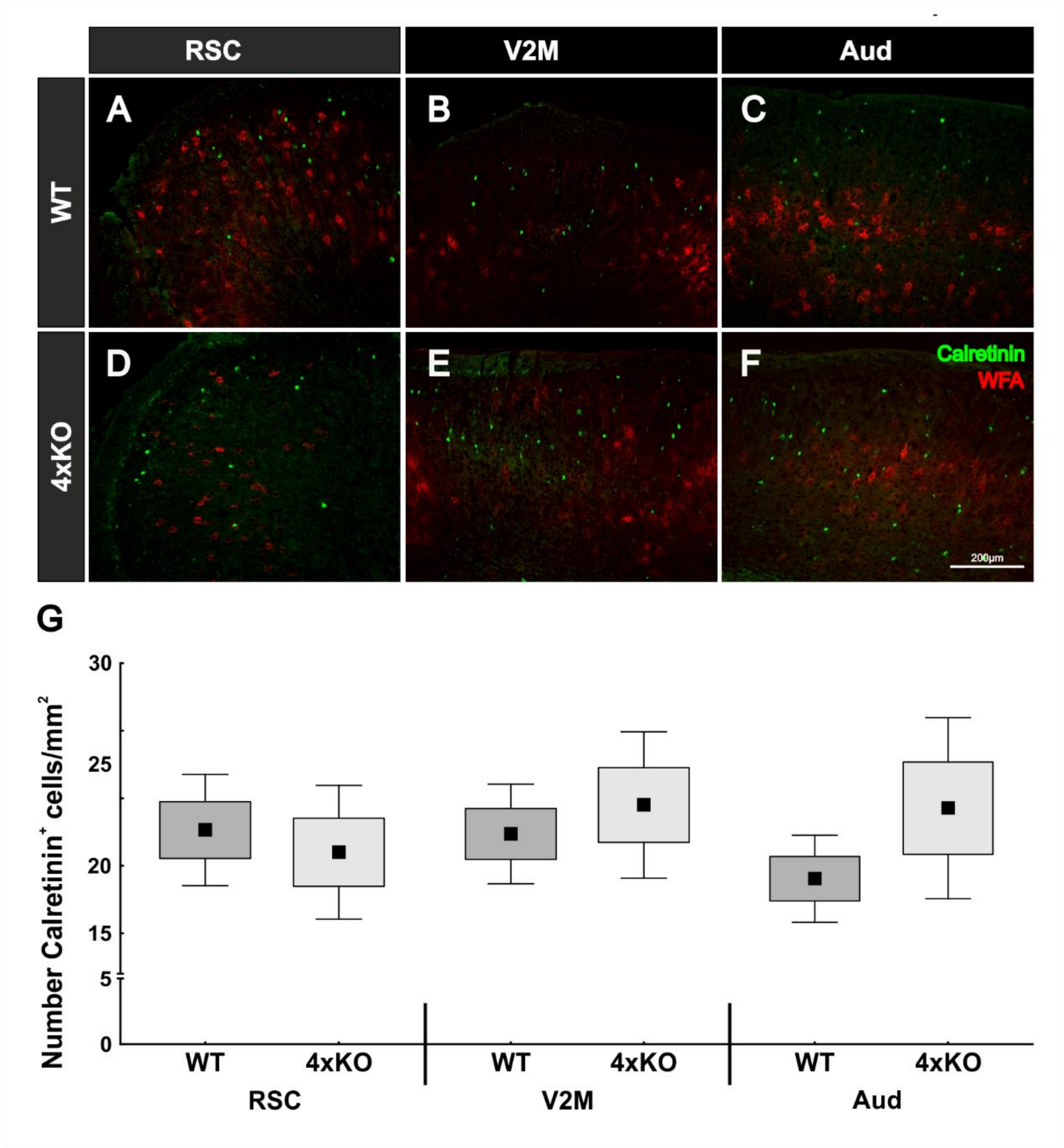
